# Quantifying (de)Mixing of Disordered Proteins in Molecular Dynamics Simulations

**DOI:** 10.64898/2025.12.12.694022

**Authors:** William Morton, Robert Vácha

## Abstract

Biomolecular condensates control the spatial and temporal organization of cellular biochemistry. Their architectures often arise from complex, multicomponent mixtures governed by weak, multivalent interactions between intrinsically disordered regions (IDRs) of proteins. However, current approaches lack generalizable metrics to determine whether IDRs will mix or segregate within condensates. Here, we present a domain decomposition method (DDM) that accurately determines phase concentrations without identifying condensate interfaces. When tested against 399 IDR simulations, the DDM recovers dense-and dilute-phase concentrations within 1% agreement of established surface-based methods. By calculating spatial variances to map three-dimensional organization, this approach successfully distinguishes homogeneous mixing from surface enrichment, core–shell architectures, and complete segregation. Applying this continuous metric to 2,068 binary mixtures of 32 distinct IDRs, we demonstrate that physicochemical similarity strongly promotes mixing. Hydrophobic IDRs mixed promiscuously across all partner chemistries, whereas charged sequences mixed in a way highly sensitive to charge complementarity, forming the most stable condensates as net charge approached zero. Compared with experimental networks, our IDR-only simulations predicted the partitioning preferences for 10 of 13 proteins, with discrepancies accurately isolating cases where colocalization strictly requires RNA or folded domains. Quantifying mixing in this way provides a robust and reproducable method for predicting IDR interactomes and understanding the composition of complex cellular condensates.

**Significance:** Biomolecular condensates are the cell’s way of organizing biomolecules into easy-to-access “containers”. Admittance can often be tuned by a protein’s intrinsically disordered regions (IDRs). However, we still lack a fundamental understanding of the features that determine whether two proteins will colocalize in the same condensate. Computational models of disordered proteins have been shown to accurately predict the condensate formation of individual IDRs. In this publication, we propose a methodology that standardizes measurement techniques and can be extended to quantify the degree of mixing between two disordered proteins. Our research shows how this relates to experimental assays, and we demonstrate the creation of disordered interaction matrices based on computational results.

## 1 Introduction

Interactions between the intrinsically disordered regions (IDRs) of proteins play a key role in driving both phase separation and partitioning into compositionally specific condensates [1–3]. The flexible nature of IDRs can create a dense network of contacts between copies of the same protein or other species[4, 5]. While some IDRs can form condensates in the absence of other proteins, they can also be distributed throughout other condensates in the cell[6, 7]. This component sharing drastically increases the number of unique condensate compositions, and similarly increases the number of interaction networks we would need to understand[8]. Within this highly interconnected landscape, disease-associated mutations could subtly alter a protein’s interaction preferences, causing it to incorrectly partition into a new condensate or disrupt the composition of its native one. These effects have been increasingly identified as a molecular hallmark of numerous human diseases, including neurodegenerative disorders and cancers[9–14]. Consequently, deciphering how IDR sequence grammars dictate condensate mixing is essential for understanding both healthy cellular organization and disease-driven dysregulation.

Assessing these IDR networks at scale can be challenging through experiments, as the human proteome is expected to contain between 14,000 and 21,000 IDRs[15, 16]. Even in high-throughput experiments that have been performed, the protein buffer, fluorescent tags, or the concentration and type of molecular crowder can all influence the interaction between two IDRs[17, 18]. Computational approaches offer a way to simulate IDR interactions without these potential confounders[19]. Despite their simplified nature, coarse-grained molecular dynamics force fields have excelled at reproducing individual IDR characteristics, such as radius of gyration, as well as bulk condensate properties such as saturation concentration and viscosity[3, 15, 20–23].

Unfortunately, current approaches to quantify IDR mixing are study-specific and thus have limitations in sequence length, maximum number of phases, or the concentration ratio of the mixture[24–27]. We are missing a generalizable method that can characterize the spatial organization of components—whether they mix homogeneously, form distinct domains, or fully segregate into separate condensates. A new characterization method could drastically improve our understanding of IDR interaction networks, leading to the development of small molecules or synthetic peptides that can selectively dissolve aberrant condensates or stabilize essential ones[28, 29].

In this manuscript, we present a methodology for both calculating an IDR’s propensity for phase separation and quantifying IDR mixing from direct coexistence simulations. Our method is designed to be simple and to improve standardization of results across studies by eliminating user-defined choices of surface thicknesses or phase boundaries. We accomplish this by adapting a method developed to characterize liquid-liquid phase transitions in simple Lennard-Jones fluids and extending it to the more complex case of multi-component biomolecular condensates[30, 31]. We test the methodology against a chemically diverse set of IDRs and use it to identify features that govern whether two IDRs mix within a coarse-grained simulation.

## 2 Results & Discussion

### 2.1 Phase Identification with Domain Decomposition

We developed and tested our domain decomposition method (DDM) to achieve two goals when analyzing direct coexistence simulations. The first is to eliminate the need to identify condensate interfaces, where the methodology can vary from study to study. Our second goal is to improve our understanding of internal condensate organization by moving from a 1- to 3-dimensional description of the dense phase.

Our DDM starts by centering the largest contiguous density in the trajectory along the extended axis of the simulation box. The volume is then split into equally sized voxels, of side length S, with fixed positions. In each voxel, the concentration is averaged over time and then fit to a kernel density estimator (KDE). Each phase in present in the system maps to one peak, the center of which marks the phase concentration. Uncertainty is then calculated by block averaging the per-frame concentration of the voxels labeled dense and dilute, where a voxel is labeled by which peak its time-averaged concentration falls under[32]. Voxels at the interface fall under neither and are excluded. An example of this process is shown in Figure 1A, with a snapshot from an example trajectory, the instantaneous concentration in each voxel during the last timestep, and the time-averaged probability density.

**Figure 1.**
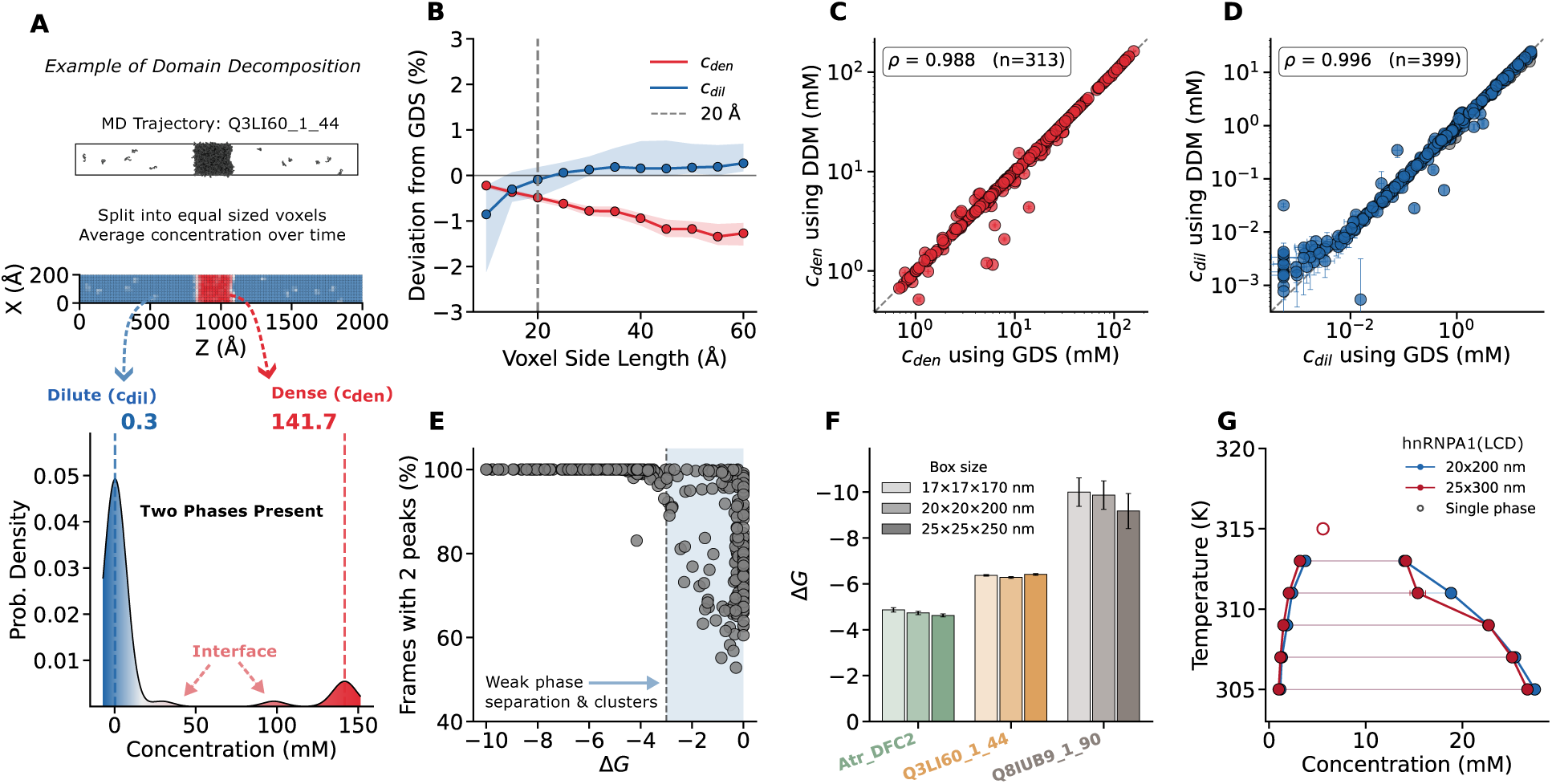
Validation of the domain decomposition method (DDM). **(A)** DDM applied to a direct coexistence simulation of UniProt:Q3LI60, residues 1–44. Top: concentration of each voxel in the final frame, averaged over Y for display. Voxels divide the cell into equally sized prisms in three dimensions. Bottom: probability density of the time-averaged voxel concentrations. The two peaks give the dilute / dense concentrations, and the shoulder below the dense peak comes from interfacial voxels. **(B)** Median deviation of DDM from GDS concentrations as a function of voxel side length, with 95% bootstrap confidence intervals (shading). Both phases agree with the GDS to within ∼1% at every voxel size. *c_den_* drifts increasingly negative as larger voxels mix interfacial residues into the dense phase, while *c_dil_* is essentially size-independent above 20 Å. The dashed line marks the 20 Å voxel used throughout. **(C)** Dense- and **(D)** dilute-phase concentrations from the DDM (*S* = 20 Å) versus the Gibbs dividing surface method. Error bars on both axes are block-averaging uncertainties; the dashed line is *y* = *x*. Grey points in (D) are systems in which no dense phase was resolved, so *c_dil_* is the mean concentration over all voxels. **(E)** Fraction of frames in which a KDE of the instantaneous voxel concentrations resolves two peaks, plotted against Δ*G*. Weak phase separators (Δ*G > −*3 *k_B_T*, shaded) resolve two peaks in fewer frames because *c_dil_* and *c_den_* are too close to separate. **(F)** Δ*G* for three IDRs in three box sizes at constant residue density. Agreement across box sizes indicates the measured concentrations are free of finite-size artifacts. Error bars are block-averaging uncertainties. **(G)** Phase diagram of the hnRNPA1 low-complexity domain from DDM concentrations, in two box geometries at equal residue density. Filled points are coexisting dilute and dense concentrations joined by tie lines; open points are single-phase state points. The two geometries give the same coexistence curve.

First, we show that the DDM accurately quantifies the concentration of the dense and dilute phases ( *c_den_* & *c_dil_*) in simulations of IDRs. We compare our measured concentrations to those from von Bülow et al., which are calculated by fitting a Gibbs dividing surface (GDS) to the one-dimensional density profile, and measuring t he average concentrations on both sides of the surface[33]. Using the same trajectories from 399 simulations allows us to test a broad range of physicochemical features (Supplementary Figure S1) and multiple box sizes. In Figure 1B we show how the voxel side length, the only user-determined parameter, affects the agreement between the two methods over the range 10–60 Å. Agreement is quantified by ln(*c^DDM^ /c^GDS^*) for *c_den_* and *c_dil_* across simulations, which we convert to a percentage deviation for interpretability.

The dense phase concentration remains within 1% agreement across all voxel sizes, but larger voxels are increasingly contaminated by interfacial residues, leading to an underestimation of *c_den_*. In small voxels (10-20 Å), the average occupancy of a dilute-phase is less than 1 residue/voxel, skewing the KDE toward 0 and underestimating *c_dil_*. Larger voxel sizes (25-60 Å) tend to overestimate *c_dil_* due to the inclusion of the interfacial IDRs in voxels near the condensate surface. Despite these effects, the DDM reliably distinguishes the two phases at every voxel size tested. We choose to use *S* = 20Å for the remainder of this study, due to its excellent agreement for both phase concentrations.

In Figure 1C & D, we take a closer look at how the two methods compare with *S* = 20Å. Out of 399 simulations, only 313 had an identifiable dense phase by both methods. Of the 86 IDRs in the single phase group, 81 had a resolved dense phase by one of the two methods. However the median transfer free energy of these sequences was Δ*G* = *k_B_T* ln(*c_dil_/c_den_*) = *−*0.19*k_B_T* [34] indicating weak, if any phase separation[33]. What either method resolves in these cases is not a stable condensate but transient clusters of IDRs.

To better understand these weak phase separators, we examine how consistently two phases appear using the DDM. Within each frame, we fit a KDE to the instantaneous voxel concentrations, and calculated the percent of frames with two resolved peaks (Figure 1E). Systems with Δ*G < −*3*k_B_T* resolve two peaks in essentially every frame, while the percentage in weaker systems becomes highly variable. Crucially, the spread at Δ*G* = 0 shows that consistent identification of two phases is a poor metric for determining phase separation. In the remainder of this study, we therefore classify a system as phase separating only when Δ*G < −*3*k_B_T*.

We next focus on the low concentration region of c*_dil_*, where both methods have the largest proportional uncertainty. Strong phase separators rarely sample the dilute phase, making it difficult to quantify. Despite having similarly large error bars, the DDM generally reports a higher concentration at low c*_dil_*. This is likely because chains leave the interface without traveling far enough to be counted as dilute by the GDS before re-entering the condensate. However, since the DDM makes no assumption about interface location, we can better capture these rare events.

Finally, we asked whether the DDM’s concentrations depend on the size of the simulation cell. Three proteins spanning a range of Δ*G*, and originally simulated at 20×20×200 nm, were re-run at with the same residue density in box sizes of 17×17×170 nm and 25×25×250 nm. All three recover the same Δ*G* within uncertainty across every box (Figure 1F). As expected, the strongest phase separator, Q8IUB9, has the largest uncertainty in each of the simulations. The sampling of the dilute phase is likely enhanced at larger box sizes, which have larger interfaces but are slower to run (ns/day).

Finally, we simulated the temperature coexistence curve for the low-complexity domain of hnRNPA1 between 305 and 319 K in two box geometries with equal residue density. The same phase diagram is recovered from both geometries, with no dense phase above 313K identified in either system (Figure 1G). The disagreement in dense concentrations at 311 K stems from multiple condensates forming in the larger box size, as seen in Supplementary Figure S2. When we center the trajectory on the largest dense region, this smears the second drifting condensate and causes us to underestimate the density. This limitation is common in direct coexistence analyses and usually occurs at low Δ*G*[35].

In this section, we have shown that the DDM recovers dense- and dilute-phase concentrations in agreement with the Gibbs dividing surface method across 399 simulations spanning a wide range of sequence lengths and physicochemical properties, without requiring an interface to be located. However, we note that this validation is specifically for direct coexistence simulations of disordered proteins, for which we have a large dataset to compare with. Determining whether this method is appropriate in cubic simulations, or in simulations with structured proteins, is outside the current study. In the following sections, we will use this same methodology to investigate the internal organization of two-component condensates.

### 2.2 Two-Component Mixing and Demixing

Binary IDR mixtures can exhibit diverse spatial organizations, from complete mixing in a single condensate to full segregation. Between these extremes lies a continuum of partial mixing behaviors, stemming from differences in the preference for homo and heterotypic interactions. To systematically characterize this mixing landscape, we analyze the voxel-level residue concentrations in our DDM and present a quantitative demixing metric based on the spatial distribution of each species within the concentration space.

To start, we perform the same DDM as presented earlier, using a voxel side length of 20Å. The only difference is that we also keep track of how many residues within each voxel belong to each IDR. We then create three KDEs: one for the total residue concentration, and one for each IDR. Figure 2A shows these KDEs for a visibly well-mixed system, revealing that both IDRs have a dense peak near 13.75 residues/voxel, exactly half of the mixed dense phase concentration. The two KDEs are computed separately, so they cannot tell us whether a voxel rich in one IDR is also rich in the other. Plotting the two concentrations against each other keeps them paired, with each point representing the concentration of both IDRs in a single voxel (Figure 2B). The KDEs from Figure 2A appear along the outer edges of this plot. We intentionally use residues per voxel here rather than molarity to avoid unequal scaling caused by different chain lengths, ensuring the two species can be visualized proportionally in the plots. These values can be converted to molarity using

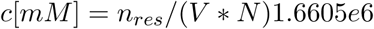

where *n_res_*is the residue count in a voxel, V the voxel volume in Å^3^, and N the number of residues per chain.

**Figure 2.**
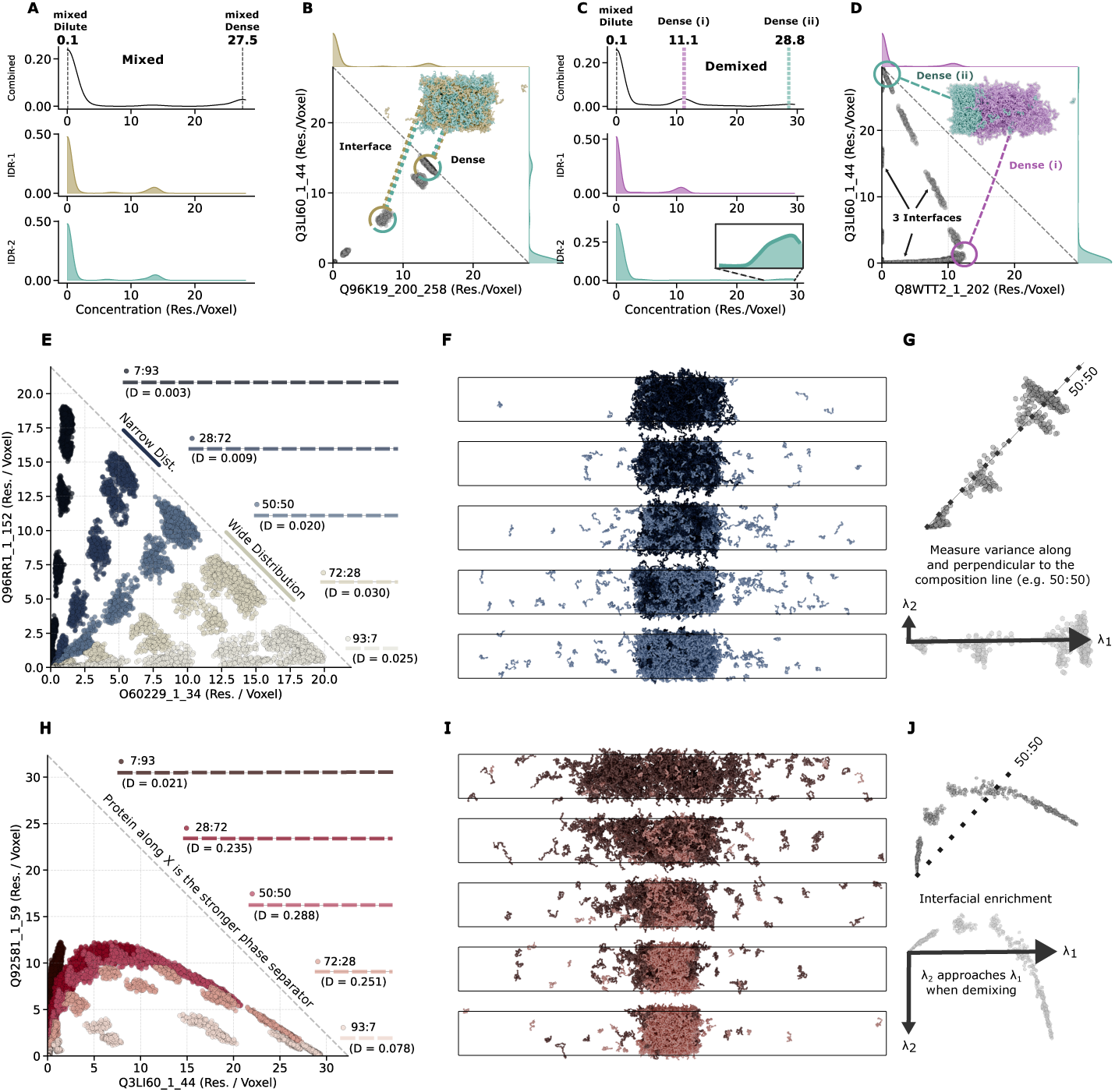
Quantifying mixing and demixing from the voxel concentration space. Panels **(A)** and **(B)** show our domain decomposition analysis for two IDRs that mix well, showing IDR-1 (Q96K19_200_258) in yellow and IDR-2 (Q3LI60_1_44) in teal. **(A)** We decompose the concentration based on the contribution from each species. Here,both IDRs have a dense peak near 13.75 residues/voxel, contributing equally to the mixed dense-phase concentration of 27.5 residues/voxel. **(B)** Each scatter point represents the average concentration of each IDR in one voxel from the direct coexistence simulations. The same probability distributions from **(A)** are shown on the outer edges of the plot. A colored snapshot from the trajectory shows IDR-1 (Q96K19_200_258) in yellow and IDR-2 (Q3LI60_1_44) in teal. The scatter points representing the surface and dense phases are circled and connected to their corresponding areas in the snapshot. Panels **(C)** and **(D)** depict our domain decomposition analysis for two IDRs that do not mix (demix), showing IDR-1 (Q8WTT2_1_202) in purple and IDR-2 (Q3LI60_1_44) in teal. **(C)** For demixed IDRs, three peaks are present in the combined probability density plot. The first and second dense phases (Dense i / ii) align with the dense concentrations of IDR-1 and IDR-2 respectively. **(D)** Plot of the voxel concentration space for a demixed system. The scatter points representing the two dense phases are circled and connected to their corresponding areas in the snapshot. Points representing the three interfaces within the system are also highlighted. **(E)** Concentrations of UniProt:O60229_1_34 (light blue) and UniProt:Q96RR1_1_152 (dark blue) in voxels from simulations with varying compositions of the two IDRs. Composition ratios are residue fractions and are shown in different colors to aid interpretation; the dashed lines mark the compositions whose final frames are shown in panel **(F)**. The demixing index *D* is shown for each sample; the largest is at a ratio of 72:28, due to the wide distribution of points within the dense phase. **(F)** Snapshots from the five compositions presented in panel **(E)**. **(G)** The variance perpendicular to the composition line (*λ*_2_) captures the width of the distribution in the dense phase and is small compared to *λ*_1_. **(H)** As in panel **(E)**, we show the concentrations of UniProt:Q3LI60_1_44 and UniProt:Q92581_1_59 in simulations with varying compositions of the two IDRs. Q3LI60 is a much stronger phase separator, with Δ*G* = *−*5.7 *k_B_ T*, compared with Δ*G* = *−*2.9 *k_B_ T* for Q92581. This imbalance and the heterotypic interaction between the IDRs enrich the surface of the dense phase in Q92581. **(I)** Snapshots from the five compositions presented in panel **(H)**. The formation of a dense phase is highly dependent on Q3LI60 (light red), while Q92581 (dark red) is localized towards the surface. **(J)** The compositional variance (*λ*_2_) is larger due to the enrichment of Q92581 at the interface. These points specifically fall roughly halfway along *λ*_1_.

When two IDRs demix into separate condensates (Fig 2C&D), distinct peaks for both dense phases may appear in the density distribution. However, the height of these peaks will be limited by the number of voxels with the dense phase present, as is the case for Dense (ii) in Figure 2C. The concentration space for this system (Fig 2D) shows three unique interfaces: one along the x-axis for the dense-to-dilute phase of Q8WTT2, one along the y-axis for the dense-to-dilute phase of Q3LI60, and one from the dense-to-dense phase where the two protein condensates meet. While visually interpreting these plots can help us understand the organization of the dense phase(s), we need a more high throughput way to quantitatively compare these mixtures.

To address this need, we developed an equation to quantify the demixing present in concentration space with a single number that we call the demixing index, D.

The demixing index is built from the composition line, which represents the global ratio of residues from the two IDRs. As an example, take the case where 50% of the total residues in the system below to each IDR (50:50). In a perfectly mixed system, every region in the dense phase, at the interface, and in the dilute phase would have a fraction of residues proportional to the global concentration. The only variance in this system, *λ*_1_, would be the total residue concentration in each voxel, varying from empty to dense.

Now let *λ*_2_ represent the variance perpendicular to *λ*_1_. When all voxels have the same composition, *λ*_2_ = 0. Our demixing index compares these two variances,

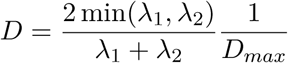

and ranges from 0, when every voxel falls on the composition line, to 1, when the two variances are equal. Taking the minimum ensures this range without assuming which of the two is larger. A normalization factor, D*_max_*, is applied to account for the reduced variance possible at unequal ratios (e.g. 90:10), and is derived in the Methods.

In Figure 2E&F, an example of a mixed system is shown at five different compositional ratios, with the total number of residues kept constant. Although each of these mixtures exhibits only one dense phase, the concentration space shows different behaviors as the fraction of O60229 (light blue) increases. Noticeably, the 28:72 ratio shows a narrow distribution of points in the dense phase, while the reciprocal ratio (72:28) shows a greater spread. This wider distribution indicates an increasing preference for homotypic over heterotypic interactions, which is captured in the demixing index reported for each system. In Figure 2G, we show the two variances for the 50:50 ratio, where *λ*_1_ characterizes the variance in density, and *λ*_2_ characterizes the variance in composition.

Figures 2H&I show a more complicated system, in which Q3LI60 (light red) is a much stronger phase separator than the Q92581 (dark red), with ΔΔ*G* = 2.8*k_B_T*. Here, the mixture ‘leans’ towards a pure dense phase of Q3LI60 at the center of the condensate while Q92581 is enriched at the interface. The variance for the (50:50) mixture of this system does not fall along the composition line, but instead arcs from one side to the other. Points with a higher concentration, towards the maximum of *λ*_1_, are those within the dense phase of the condensate and have a larger fraction of Q3LI60. Those towards the middle of *λ*_1_ are localized on the interface where Q92581 is enriched.

While other demixing metrics have been proposed, they’ve either characterized core–shell condensates, assume that each component’s concentration is radially symmetric about the condensate center, require precise identification of phase boundaries, or some combination of these[26, 27, 36]. To show that these assumptions do not apply for our MD systems, we show the one dimensional profiles of several systems with varying levels of demixing, and different condensate architectures, in Supplementary Figure S3. In the Supplementary Information, we show a comparison to a robust mixing method recently introduced by Chen & Jacobs, which agrees well at (50:50) ratios, but poorly at unequal ratios (Supplementary Figure S6)[27].

In the following sections, we create a set of diverse IDRs, and explore what physicochemical properties cause some to mix while others segregate, and how their mixing affects the thermodynamic stability of the resulting condensates.

### 2.3 Contents of Protein Set

To investigate the determinants of IDR (de)mixing, we curated a set of 32 IDRs and performed direct coexistence simulations on all 496 unique heterotypic pairs. Our selection of IDRs spanned a range of sequence compositions and propensities for phase separation. To explore the compositional space, we selected proteins from different grammar clusters[37]. These are groups of IDRs with similar sequence patterns defined by their amino acid composition (Supplementary Table I). We specifically chose IDRs within those grammars that had varying propensities for phase separation, Δ*G*. All proteins were drawn from the von Bülow et al. dataset[33], allowing us to use simulated Δ*G* values from their published trajectories. More details on the selection algorithm are provided in the Methods section.

The resulting protein set spans a wide range of physicochemical properties and is enriched in strong phase separators. Our subset’s median length (95 amino acids) is shorter than that of the source dataset but more closely reflects the median length of IDRs in the human proteome (80 amino acids) [15]. We deliberately enriched for sequences with higher mean residue hydrophobicity (*λ*^^^) and a stronger homotypic phase separation (more negative Δ*G*) to ensure a large proportion of condensate-forming mixtures. The subset adequately samples sequences with varying aromatic content, ranging from aromatic-poor to aromatic-rich. Our sampling of the charged sequence space is narrower, with the net charge per residue (NCPR) *≈* 0 where homotypic phase separation is more prevalent. The full distribution of each feature, in comparison to the original dataset, is shown in Supplementary Figure S1.

For each unique pair in our protein set, we performed simulations at equal residue compositions (50:50) with *∼* 36, 000 amino acids in total. In a simulation where one protein has 50 residues and the second has 100, this would mean 360 copies of the former and 180 copies of the latter. Additional concentration ratios were simulated (e.g., 25:75 and 90:10) with the same number of amino acids, where protein lengths permitted. In total, we performed 2068 simulations, all of which started in a completely demixed state.

### 2.4 Chemical Grammar Influences Mixing Behavior

Sequence features have previously been used to group IDRs into self-similar families and to predict whether an individual protein will phase separate. For example, high fractions of aromatic residues or well-distributed charge blocks generally favor condensate formation[38, 39]. It follows that modifications to these features will alter a protein’s capacity to mix with partners in a condensate, but simulating each variant can be redundant and computationally costly. Here we asked whether the same features and chemical characteristics can also be used to intuit which pairs of IDRs will mix *in silico*.

Each IDR was assigned the parent chemical character of its GIN cluster — charged (C), polar (P), or hydrophobic (H) — and every pair placed into one of six interaction categories (Supplementary Table S1). Of the categories shown in Figure 3A, hydrophobic-hydrophobic pairs have the lowest demixing indices (median *D* = 0.015, n = 113), which we attribute to the non-specific nature of hydrophobic contacts. This is consistent with recent experimental work, in which the average fraction of aromatic residues (F, Y and W) between two IDRs correlated positively with their miscibility[40]. One hydrophobic cluster in particular, well-mixed hydrophobics enriched in methionine (GIN cluster 10), mixes readily not only within the H-H category but with every partner chemistry we tested (median *D* = 0.023, n = 436). IDRs of this type are common in cancer-associated proteins, where they form condensates that localize transcriptional machinery to DNA[14, 37, 41]. Mutations that alter the phase separation of these proteins could therefore have larger knock-on effects that stem from this promiscuity. A mutation stabilizing one of these condensates could also prolong the retention of its mixing partners, a proposed mechanism for the dysregulated transcriptional machinery observed in leukemia[42]. While our dataset is limited, these results suggest that the mixing of nuclear IDRs could be further explored *in silico*.

**Figure 3.**
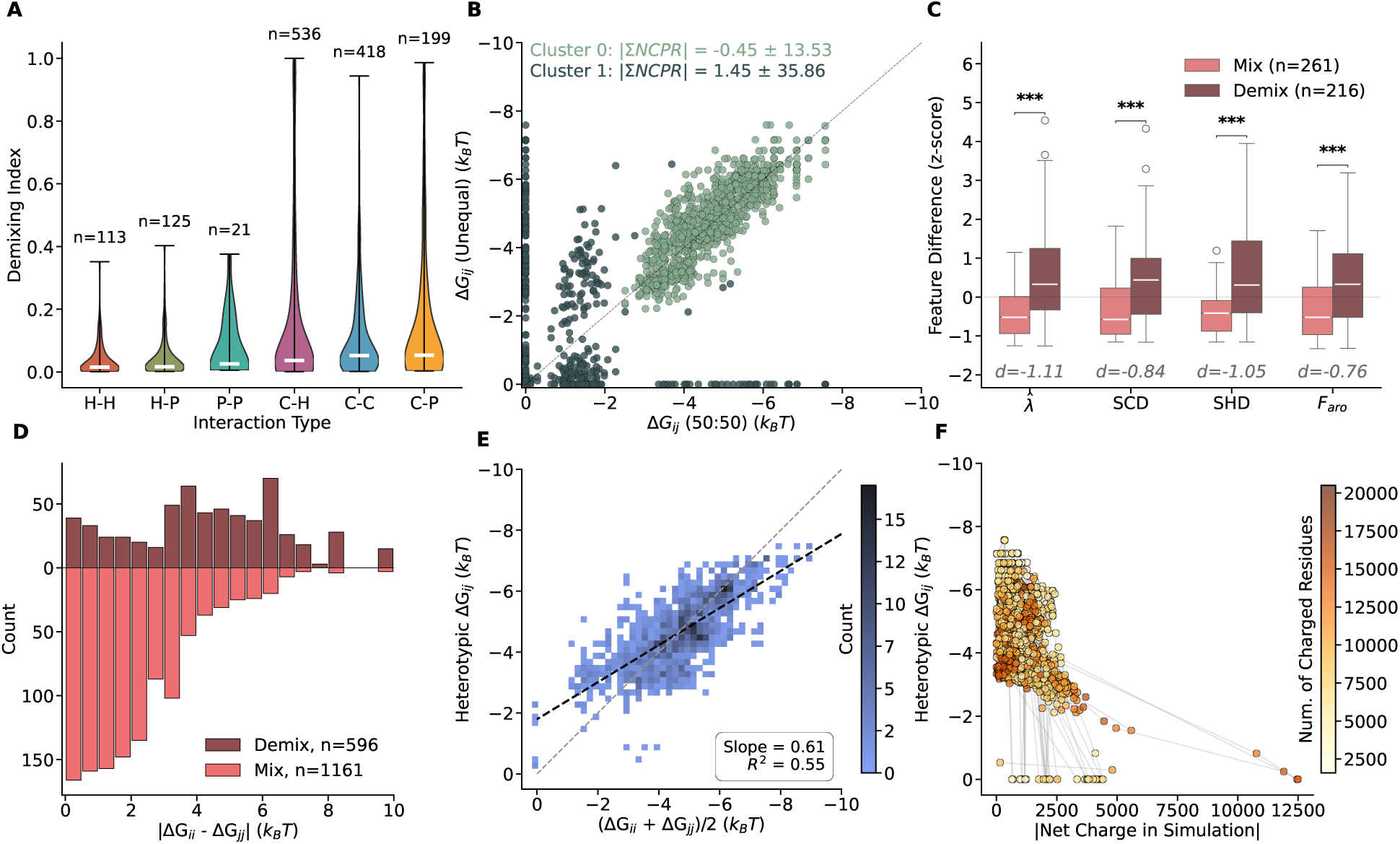
Chemical and Physical Properties of Mixed Condensates. **(A)** Distribution of demixing indices across interaction categories defined by the dominant chemical character of each IDR: C = charged, P = polar, H = hydrophobic. All condensate-forming simulations are included at every concentration ratio. **(B)** Comparison of heterotypic phase separation strength (Δ*G_ij_*) at (50:50) versus unequal compositions. We split the data into two clusters using a Gaussian Mixture Model on the Δ*G* values. The sum of net charge per residue (|ΣNCPR|) is shown for the two groups. Neutral mixtures show consistent stability across compositions (lying on the diagonal), while charged mixtures deviate significantly. **(C)** Feature difference analysis for mixed ( blue) versus demixed (pink) pairs at (50:50). The plot shows the difference in z-scores for mean residue stickiness (*λ*^^^), sequence charge decoration (SCD), sequence hydropathy decoration (SHD), and fraction of aromatic residues (*F_aro_*). Mixed pairs exhibit significantly smaller differences in these physicochemical properties (*** *p <* 10*^−^*^6^), with strong effect sizes (Cohen’s *d*) annotated below each feature. **(D)** Distribution of the difference in homotypic phase-separation propensity (|Δ*G_ii_ −*Δ*G_jj_*|) across all compositions, using the same mixing criteria as panel C. Well-mixed condensates predominantly form when IDRs have similar phase-separation propensities (difference *<* 4 *k_B_T*). **(E)** Heterotypic Δ*G_ij_* versus the average of the two homotypic values ((Δ*G_ii_*+Δ*G_jj_*)*/*2). The dashed gray line indicates Δ*G_ij_* equal to the average. The linear fit (black dashed line) shows that the average homotypic Δ*G* correlates with heterotypic stability, though charged sequences often exhibit stronger heterotypic interactions due to improved charge balancing, which can be seen in Supplementary Figure S5. **(F)** Heterotypic Δ*G_ij_* as a function of the total number of unbalanced charges in the simulation. Lines connect different compositions of the same protein pair. As net charge approaches zero, condensate stability (more negative Δ*G_ij_*) generally increases. Point shading indicates total charged residues, revealing that the strongest phase separators are not necessarily the most charged, suggesting contributions from other interactions such as aromatic contacts.

In contrast to the promiscuity of hydrophobic sequences, we expected mixing with charged sequences to require some degree of charge complementarity. While charge–charge pairs span a broad range of *D*, they did not fully demix. We attribute this to a conflict between the requirements for demixing and those for charge-driven phase separation. Complete demixing typically requires at least one partner to phase separate strongly on its own, thereby excluding the other. For a charge-dominated IDR, however, strong homotypic separation requires a balance of positive and negative residues, meaning a charged partner could be partially miscible. However, it’s unclear how relevant this would be to cellular condensates, which have several mechanisms to control the electrochemical equilibria in condensates[43].

We next investigated the compositional dependence on condensate formation, which we expected to be most affected by sequence pairs with more charges. Figure 3B compares the strength of phase separation at (50:50) compositions to other ratios. After identifying a natural split in the scattered points, we used a Gaussian mixture model to create two clusters based on the Δ*G* values alone. We then measured the summed net charge per residue (|Σ*NCPR|*) of each point in the two groups. Pairs along the diagonal have almost no charged residues, suggesting that changes in composition cause the mixed system to move between the Δ*G* of the two individual IDRs. Pairs in the second group, with |Σ*NCPR|* = 0.15, straddle the diagonal and sit along the axes, indicating that condensate formation is dependent on charge balance, rather than homotypic Δ*G* values.

The smallest grouping in this study is that formed by polar–polar pairs. IDRs in the polar GIN grammars are predominantly hydrophilic and rarely phase separate alone. As a result, they are underrepresented within our sample set. Only three of the five IDRs in the polar group undergo homotypic phase separation, which we attribute to their high aromatic content. Every P–P condensate we observe involves at least one of these three. In the single pair where neither partner phase separates alone, no composition produced a dense phase.

The low demixing seen for chemically self-similar pairs, most clearly for H-H, prompted us to ask whether IDRs with similar physicochemical properties preferentially co-condense. In this analysis, we split (50:50) mixtures into two groups. The first group contains those that mix well (*D <* 0.2) and phase separate (Δ*G_ij_ < −*3*k_B_T*). The second group contains those that do not mix (*D ≥* 0.2) but have one component that can form a homotypic dense phase, representing genuine segregation of two species rather than the trivial case in which neither partner condenses at all. For each pair, we looked at the difference in the following features: mean residue stickiness (*λ*^^^), sequence charge decoration (SCD), sequence hydropathy decoration (SHD), and fraction of aromatic residues (*F_aro_*)[44]. A two-sample t-test between the mixed and demixed groups (*n_mix_* = 261*, n_demix_* = 216) shows a significant difference (*p <* 10*^−^*^6^) for each feature. Effect sizes were large in every case, with Cohen’s d between -0.76 and -1.11. In every case the mixed group showed smaller feature differences than the demixed group, at least across the sequence space sampled here, suggesting that physicochemically similar IDRs are more miscible.

We emphasize that the patterns reported in this section are descriptive of this dataset rather than predictive of IDRs as a whole. Explicit simulations are still necessary to quantify heterotypic driving forces until larger datasets can support a general predictor. Having established which physicochemical properties cause our pairs to mix, we next investigate whether these same properties influence condensate stability.

### 2.5 Stability of Mixed Condensates

Two IDRs may mix homogeneously yet form only a marginally stable condensate. We therefore asked how the homotypic Δ*G* determines the stability of well-mixed two-component condensates. To do so, we analyze the heterotypic transfer free energy (Δ*G_ij_*) for well-mixed systems (*D <* 0.2) at every composition, analogous to how we calculate the homotypic Δ*G* using the DDM.

We first hypothesized that a stable heterotypic condensate requires the two IDRs to have comparable self-interaction strengths, since a strong phase separator may exclude a weaker interacting partner. Figure 3D shows that mixing predominantly occurs when the two IDRs have similar self-interaction strengths, with differences of less than 4*k_B_T*. Larger disparities in homotypic Δ*G* tend to demix as the stronger phase separator dominates the dense phase while the weaker partner is partially or fully excluded. These patterns are dominated by mixtures with a charged sequence. Hydrophobic and polar sequences mix well even when paired with dissimilar partners. A breakdown of the contribution from each chemical interaction group can be seen in Supplementary Figure S4.

If stability is governed by how similar the two partners are, then the average homotypic transfer free energy 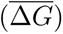 should be a reasonable predictor of Δ*G_ij_* in mixed systems. Figure 3E shows that this holds for most pairs, with a high density of points clustered around the parity line. This pattern holds best for hydrophobic pairs (H-H) which have a slope of 0.67 (Supplementary Figure S5). Unsurprisingly, the worst agreement came from pairs in which both IDRs are categorized by charge (C-C, *R*^2^ = 0.49), likely due to the need for charge balancing.

Figure 3F plots Δ*G_ij_* against the absolute net charge in the simulation, where each protein pair is represented by connected points spanning different concentration ratios. As expected, when the net charge in the system approaches zero, the strength of phase separation increases. However, from this graph we can see that the system can still phase separate even when 7% of the charges are left unbalance. This suggests that while condensate neutrality contributes to heterotypic stability, other interactions, such as the fraction of aromatic and hydrophobic residues, provide a larger driving force for condensate formation.

Taken together, these results suggest that heterotypic condensate stability follows simple patterns. Similarity in homotypic ΔG promotes mixing; charge complementarity can enhance phase separation; and nonspecific hydrophobic interactions provide a baseline driving force insensitive to partner identity.

### 2.6 Comparison to Experiments

Finally, we focus on how the methodology presented here can complement experimental studies. While the phase separation of individual IDRs has been compared elsewhere, there are few experiments where we can compare the quantitative degree of mixing. In the following section, we use our methodology to understand three experimental datasets, each of which characterized mixing in different ways.

We start by creating mixtures of fused-in-sarcoma (FUS) and hnRNPA1 (A1) (FUS-A1_WT_ - red) at three concentration ratios, to compare with the experimental studies done by Farag et al., as seen in Figure 4A&B[24]. Experimentally, the curvature of the dilute arm of the binodal corresponds to the balance of homotypic vs. heterotypic interactions between two IDRs[17, 47]. Concave dilute arms are associated with strong heterotypic interactions, while convex curves indicate a preference for homotypic interactions. For FUS-A1_WT_ the dilute arm is concave, indicating dominant heterotypic interactions. When Farag et al. substituted 12 Asp residues into the A1 (A1*_p_*_12_*_D_* - blue), the arm became convex, yet the two IDRs remained colocalized, as observed by fluorescence microscopy.

**Figure 4.**
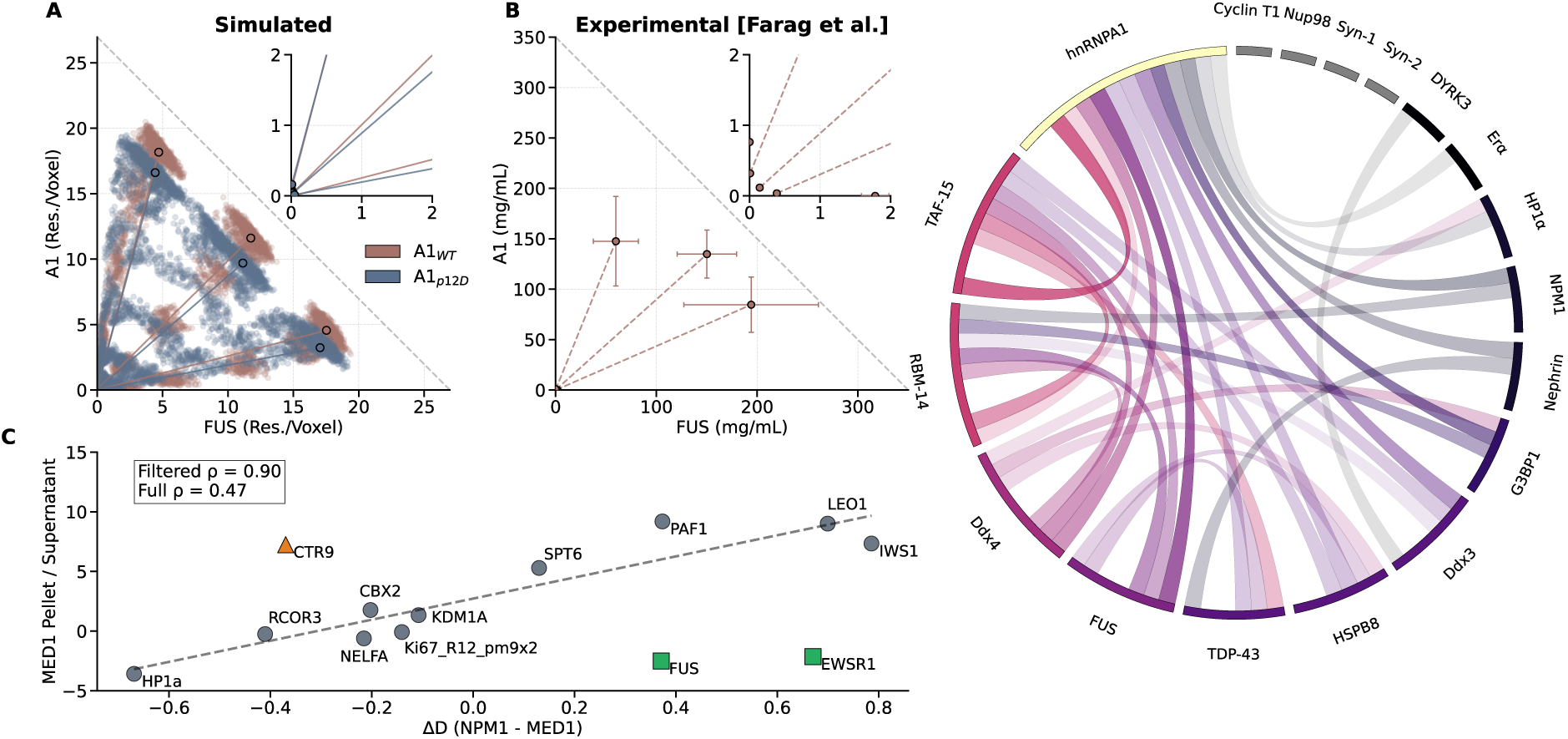
Complementing Experimental Results Using Simulated Demixing Values. **(A)** Results from simulating a mixture of the low-complexity domains of hnRNPA1 (A1_WT_) and Fused-in-sarcoma (FUS), shown in red. Simulated results from the FUS mixture with a mutated version of A1 (A1*_p_*_12_*_D_*) are shown in blue. Tie lines, connecting the dense and dilute concentrations, for both systems are calculated using the DDM and overlaid on the scattered points. The inset of the plot shows the dilute end of the tie lines, where the mixture with A1_WT_ is convex, while the mixture with A1*_p_*_12_*_D_* is concave. **(B)** Experimentally determined phase map of A1_WT_ and FUS at 16°C, reproduced here using the data published by Farag et al. [24]. **(C)** Comparison of simulated demixing preferences with experimental partitioning data from Lyons et al.[45]. The X-axis shows the difference in demixing index between NPM1 and MED1 for each protein; positive values indicate a mixing preference for MED1, negative values for NPM1. Ten of thirteen proteins show demixing preferences consistent with their experimental pellet/supernatant distribution. FUS and EWSR1 are shown in green because they both exhibit high demixing with NPM1 and MED1 and low demixing with each other. CTR9 is shown in orange because it has been shown experimentally to phase separate with MED1 in the presence of RNA, which is not present in these simulations. The Spearman’s p is shown for all 13 data points (Full) and for the 10 not considered outliers (Filtered). **(D)** IDR interaction map for the 18 sequences studied by Gilat et al., where each unique pair was simulated at (50:50) compositions[46]. The color of each chord is related to the color of the partner IDR. The thickness is related to the demixing index of the connected pair. Thicker lines denote lower demixing indices. The transparency of the chord is related to the relative strength of phase separation (Δ*G_ij_*). No connection is shown between pairs that did not form a dense phase, nor those with (*D >* 0.2).

Our simulation results similarly show colocalization, weaker phase separation, and a higher D for the mutated sequence. However, the dilute arm of the simulated mixtures shows the opposite behavior to that in the experimental study. As previously stated, the dilute-phase concentration is often undersampled in MD simulations, whereas the dense phase converges quickly. The opposite is true for experimental studies, where the dilute phase concentration is easily measured by centrifuging the dense phase out of solution and then measuring the protein concentration in the supernatant. Therefore, we suggest that the demixing index and the concentration of the dense phase are better metrics for comparison with experimental observables.

Next, we turn to the study by Lyons et al., which co-purified the IDR of the Mediator-1 complex (MED1), a transcription regulator, with soluble nuclear extract[45]. The MED1 condensate and everything partitioning into it were separated by centrifugation (pellet), while the remainder stayed in the supernatant. The authors noted that proteins excluded from MED1 tended instead to partition with NPM1[48]. We simulated the mixing of 13 of these proteins with both MED1 and NPM1 and compared their demixing preferences against the experimental pellet-to-supernatant ratio (Figure 4C). Ten of the 13 show a clear preference for one partner that matches the experimental fractionation (Spearman’s *ρ*_10_= 0.90; Spearman’s ρ_13_= 0.47).

We highlight these three outliers because their poor correlation may be attributable to differences between our experimental and simulated systems. FUS and EWSR1 (green) have similar physicochemical properties, mix well with one another, and demix strongly from both MED1 and NPM1 (Table 1). Their enrichment in the supernatant is therefore unlikely to be due to affinity for either scaffold. In the NPM1-induced lysate granules of Freibaum et al., both proteins were enriched well below the population median: 5.9-fold for FUS and 20-fold for EWSR1, against a median of 40-fold across the 1,116 proteins detected[48]. Both are canonical stress-granule components rather than nucleolar ones, and their partitioning in cells depends on RNA and on folded domains that our IDR-only simulations omit. CTR9 (orange), on the other hand, is well known for its colocalization with MED1 in both experiments and coarse-grained simulations [49–51]. However, in every case, RNA is required to drive the two into a condensate, and no RNA was present here. The juxtaposition between the excellent agreement and the outliers makes simulated results difficult to interpret. MD can likely capture how IDRs drive condensate formation and partitioning, unless an additional component, such as folded domains or RNA, is required. Additionally, partitioning in the experiment reflects a multicomponent network in which each protein competes with hundreds of others, whereas our simulations test the interaction between two IDRs at a time.

**Table 1.** Demixing indices (D) from (50:50) direct coexistence simulations. MED1 and NPM1 failed to form a condensate when mixed.

|  | FUS | EWSR1 | MED1 |
| --- | --- | --- | --- |
| EWSR1 | 0.08 |  |  |
| MED1 | 0.62 | 0.32 |  |
| NPM1 | 0.99 | 0.98 | NA |

For our last comparison, we look at a network of interactions, using the 18 IDRs studied by Gilat et al.[46]. In that study, interactions between IDRs were analyzed by connecting them to fluorescent reporters and oligomerizing domains. Colocalization between two constructs, with unique IDRs and fluorescent reporters, was measured via confocal microscopy and used as a measure of mixing. Here, we simulate all 18 unique pairs using a (50:50) composition, following the same methodology presented earlier. Figure 4D presents the resulting interaction map as a chord diagram: a chord is drawn between two IDRs only when the simulation produced a dense phase with low demixing (D < 0.2), so the map reports which IDRs are expected to co-partition on the basis of their disordered regions alone.

Six of the seven top mixers (hnRNPA1, TAF-15, FUS, RBM-14, HSPB8, & TDP-43) demonstrated strong phase separation in the homotypic experimental constructs and exhibited similar promiscuity when mixed with various partners in vitro. In the experimental study, TDP-43 did not mix with FUS, hnRNPA1, HSP88, RBM-14, or TAF15. However, this discrepancy is due to the *α*-helical nature of residues 320-343, which were kept disordered in our study and have been noted to greatly increase homotypic interactions when folded[52]. The removal of this region caused TDP-43 to mix promiscuously with each of the aforementioned proteins, suggesting that specific condensate partitioning may be fine-tuned by folded regions, or other components such as nucleic acids and post-translational modifications.

The only other IDR that exhibited demixing in the experimental results was DYRK3, which is a weak phase separator, described by the GIN category 0:Blocks of G and Polar residues. Following the trends established above for polar sequences, DYRK3 lacks both the hydrophobic content that drives non-specific mixing and the charged residues that enable complementary heterotypic interactions, leaving no strong driving force for mixing with the other IDRs.

The IDRs in Figure 4 are integral components of distinct cellular bodies, including paraspeckles, stress granules, and transcriptional condensates[24, 45, 46]. By successfully reconstructing their mixing preferences *in silico*, we demonstrate the potential of this methodology to predict IDR interactomes. Scaling this approach would allow for the construction of interaction matrices, providing a clearer understanding of the IDR role in cellular organization. However, this section has also exemplified the limitations in this method. Other interactions, which may be necessary for promoting (de)mixing, are not captured in these models. Therefore, it’s important that this work is complemented by other models, predictors, or experimental validations.

## 3 Conclusions

We have introduced a new methodology for analyzing the phase separation of disordered proteins and their mixing, *in silico*. Our domain decomposition method accurately determines the concentrations of both the dense and dilute phases in single-component systems, without requiring identification of the condensate surface. In two-component systems, we have shown that the mixing of disordered proteins can be numerically characterized at both (50:50) and unequal concentration ratios.

By analyzing 2,068 binary mixtures, we found that, within our dataset, physicochemical similarity is a good indicator for IDR miscibility. We observed that hydrophobic sequences mix promiscuously across partner chemistries, whereas charged sequences are sensitive to the complementarity of their partner’s charge. Furthermore, we observed that the strength of phase separation in heterotypic condensates largely follows the average of homotypic values, but charged sequences can exceed this average by mixing with charge-complementary species.

Finally, we used this methodology to explain the protein organization seen in experimental studies. By isolating the disordered proteins in simulations, we can identify whether colocalization between two sequences is due to affinity between the disordered regions or to additional factors such as RNA or folded domains. Our demonstration shows that simulating larger sets of proteins could help to identify condensate compositions, potential dysregulators, and ideal targeting candidates for condensate modulation.

## 4 Limitations

We validated the domain decomposition method only for direct coexistence simulations of disordered proteins represented by a single bead per amino acid, in slab geometries. Whether the approach is appropriate in cubic boxes or in systems containing folded domains remains untested. As with other methods, the concentrations are time-averaged, which requires the dense slab to remain centered throughout the trajectory. When two or more condensates coexist and diffuse independently, the secondary condensate is smeared across the averaged map and the dense concentration is underestimated. Dense phases spanning only a few voxels contribute a correspondingly small peak to the kernel density estimate, which may overlap with interfacial voxels flanking it. For weakly associating sequences, transient clusters produce a second maximum in the kernel density estimate that does not correspond to a stable condensate. The two maxima in these systems are separated by less than the width of either, so what the fit resolves is a shoulder rather than a distinct phase, and the resulting Δ*G* carries little meaning. This is why we classify a system as phase separating only when Δ*G < −*3*k_B_T*. Finally, the demixing index is only appropriate for simulations with two disordered proteins. Systems with an IDR and RNA or folded domains require further validation, which was outside the scope of this study.

## Materials and Methods

### Protein Dataset

The proteins for our study were selected from those previously simulated with CALVADOS2 to develop a phase-separation predictor[33]. This dataset contains the IDR sequences of individual proteins, calculated sequence metrics (e.g., the fraction of aromatic residues), and the measured concentrations of the dense (c*_den_*) and dilute (c*_dil_*) phases from direct coexistence simulations. The advantage of using a subset of these proteins is that their homotypic transfer free energies, Δ*G* = *k_B_T* ln(*c_dil_/c_den_*), have already been determined by simulation.

We restricted the pool to IDRs with sequence lengths *N <* 250 residues, which allowed us to use the same simulation box for all systems at a reasonable computational cost. Proteins were categorised using the 30 GIN (Grammars Inferred using NARDINI+) clusters defined by Ruff et al.[37], which group IDRs by “molecular grammar”, which include features like amino acid composition and the patterning of charged or hydrophobic residues.

Within each GIN cluster, proteins were binned by homotypic Δ*G*, which ranges from −10 *k_B_T* (strong phase separation) to 0 k*_B_*T (no phase separation). GIN clusters containing at least one protein with Δ*G < −*5 *k_B_T* were treated as phase-separating clusters; from each we selected the shortest sequence falling in each of four bins, [−10, −8), [−8, −6), [−6, −4) and [−4, −2) *k_B_T*, skipping any bin with no member. For the remaining clusters, in which all members showed weaker self-association (Δ*G ≥ −*5 *k_B_T*), we drew one sequence at random from those with fewer than 100 residues so that the group was still represented. Clusters where no IDR had previously been simulated were not included. This yielded a set of 32 IDRs spanning distinct GIN clusters (Supplementary Table S1).

The 399 single-component trajectories used to validate the method in Figure 1 were taken directly from von Bülow et al.[33] and re-analysed without re-running; their box geometries are those of the source study, in which the cell was scaled with chain length. The three-box comparison (Figure 1F) and the hnRNPA1 temperature series (Figure 1G) were run for this work using the protocol below.

### Concentration Ratios of Simulated Mixtures

For each unique protein pair, we simulated up to five compositional ratios at a fixed total residue count. Reference chain numbers were first set from a 50/50 split of 36,000 residues, giving a pair-specific target total. For each of five target residue fractions of the first protein (0.9, 0.75, 0.5, 0.25, 0.1) we then searched integer chain numbers within *±*10 of the ideal residue count and retained the candidate closest to *n^i^*_1_*^deal^* for which the remainder was divisible by the length of the second protein. Candidates were rejected if either species exceeded 95% of the residues, and each value of *n*_1_ was used at most once, so that pairs whose lengths admit fewer exact solutions simply return fewer compositions. Every accepted composition therefore contains the same number of residues, and the ratios quoted throughout are residue fractions rather than chain or mole fractions.

For example, if IDR-1 has 40 residues and IDR-2 has 120, a target fraction of 0.75 gives 675 copies of IDR-1 and 75 copies of IDR-2, for 36,000 residues in total at a 75:25 residue ratio.

This approach yielded 2068 protein combinations with varying compositions. A full CSV of all simulated pairs can be found on the GitHub repository (see *Software and Libraries*).

### Simulation Protocol

All molecular dynamics was performed in the *NVT* ensemble with OpenMM 8.2[53] on CUDA in single precision, using a LangevinMiddleIntegrator with a friction coefficient of 1 ps*^−^*^1^. Production simulations used a slab box of 20 *×* 20 *×* 200 nm^3^ and a 10 fs time step. We used the CALVADOS2 forcefield, the details of which are reproduced in the Supplementary Information[23].

Chains were initialized so that the two IDR species were completely demixed, and then simulated for 4.5 *µ*s at 293 K, with coordinates written every 1 *×* 10^5^ steps (1 ns) to give 4500 frames. At this point, we checked to see if a condensate had formed, and those with a dense phase present were simulated for an additional 1.5 *µ*s. During analysis, the first 1 *µ*s was discarded as equilibration.

### Domain Decomposition Method

The simulation cell was partitioned into voxels by dividing each dimension *L_i_* (*i ∈ {x, y, z}*) by an ideal voxel side length *S* and rounding to the nearest integer, *λ_i_* = max(round(*L_i_/S*), 1); actual voxel dimensions are *s_i_* = *L_i_/λ_i_*, so voxels may be slightly anisotropic depending on box geometry. All reported results use *S* = 20 Å; the only exception is the convergence analysis of Figure 1B, which sweeps *S* from 10 to 60 Å.

Each frame was first centered along the elongated axis. The *z* coordinates of all beads were histogrammed in 1 Å bins, and the box was translated so that the most populated bin lay at the center. The coordinates are then wrapped, and the largest contiguous run of occupied bins was then identified. Finally, the box was translated again so that its density-weighted center lay at the box center[33]. Beads were assigned to voxels by integer division of their wrapped coordinates (clipped to the last voxel along each dimension), and, for binary mixtures, counted separately for each species.

A single time-averaged voxel map was accumulated as a running mean over every frame of the production window. Phase concentrations were then read from a Gaussian kernel density estimate of this map (scipy.stats.gaussian_kde, Scott’s rule), evaluated on a 4096-point grid spanning the observed concentration range padded by 5%, with local maxima located using scipy.signal.find_peaks. The dense phase, c*_den_*, is the tallest peak after excluding the lowest-lying one; a map with fewer than two peaks yields no dense phase.

To best capture the large order of magnitude spread of *c_dil_*, we created a second KDE evaluated using ln(*c*), restricted to voxels with a nonzero time-averaged concentration (i.e. visited at least once over the trajectory). *c_dil_* was taken as the position of the lowest maximum and exponentiated. For binary mixtures, where concentrations are expressed in residues per voxel and the two peaks are well separated on a linear axis, both *c_dil_* and *c_den_* come from the single linear KDE described above, with no voxel excluded. Each species’ own dilute and dense concentration was obtained the same way, from a linear KDE of that species’ own time-averaged voxel map rather than the total, so that a demixed pair’s two condensates are each characterised by their dominant species rather than by whichever is taller in the combined map.

A voxel map was treated as resolving two phases, and c*_den_* reported, only when at least 5% of voxels sat at or above c*_den_*). Otherwise c*_dil_* is calculated as the average concentration across all voxels and no dense value is defined.

Uncertainties were obtained by block averaging the per-frame concentration of of voxels. Each voxel of the time-averaged map was labeled dense or dilute when its value fell within the full width at half maximum of the corresponding peak. Voxels falling in neither window, i.e. interfacial voxels, were discarded. The mean concentration over each labelled set was then computed frame by frame, and the standard error of the resulting time series estimated by blocking[32].

Transfer free energies for single-component systems were computed as Δ*G* = *k_B_T* ln(*c_dil_/c_den_*) and clamped at a lower bound of *−*10 *k_B_T*, so systems whose dilute phase is too sparsely sampled to resolve accumulate at this value. A system is treated as phase separating throughout this study only when Δ*G* < *−*3 *k_B_T* (see discussion on Figure 1E).

### Demixing Index

To quantify the spatial organization of a binary mixture we analyzed the distribution of voxel compositions in the two-dimensional concentration space (c_1_, c_2_), using the trajectory-averaged voxel map.

The global composition is defined by the residue fractions of the two species, *ϕ*_1_ = *M*_1_*/*(*M*_1_+*M*_2_) and *ϕ*_2_ = 1 *− ϕ*_1_, where *M_k_* is the total number of residues of species *k* in the simulation. The composition line is the ray through the origin along

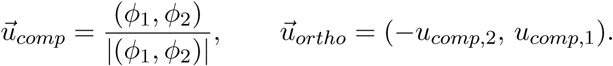

Voxel compositions (*p⃗*) were projected onto this fixed basis, giving the variance along the composition line and perpendicular to it,

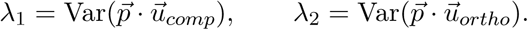

*λ*_1_ measures the spread from the dilute phase near the origin out to the dense phase and therefore reports the strength of phase separation; *λ*_2_ measures how far local composition strays from the global ratio. The demixing index is

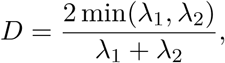

which is 0 when every voxel lies on the composition line and 1 when the two variances are equal.

We normalize D based on the composition in the simulation.

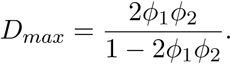

A discussion on the derivation of this factor can be found in the Supplementary Information. We report the composition normalized D/D*_max_* throughout, and refer to it as D in the text.

## Software and Libraries

- **MD simulations:** OpenMM 8.2[53]
- **Trajectory analysis:** MDAnalysis 2.9.0
- **Scientific computing:** NumPy 1.26.4, SciPy 1.13.0
- **Block averaging:** BLOCKING[32], https://github.com/fpesceKU/BLOCKING
- **Machine learning:** scikit-learn 1.5.2
- **Data manipulation:** Pandas 2.1.1
- **Visualization:** Matplotlib 3.9.3
- **Rendering:** Blender 4.2.1

## Data, Materials, and Software Availability

All analysis code, including the demixing index calculation and the full set of simulated protein-pair compositions, is available at https://github.com/shakespearemorton/IDRMixing.

## Acknowledgments

Several people helped evaluate this work, ensuring it would be useful and interpretable for broader audiences. We thank Francesco Luca Falginella for discussions and reading the full first draft; Soren von Bülow for taking the time to meet and talk about the findings; Salomé Bodet-Lefevre for applying an experimental eye to molecular dynamics figures; and Erin Spearing for letting me talk about this work incessantly, and providing experimental insights. This project has received funding from the European Union’s Horizon Europe research and innovation programme under the Marie Sklodowska-Curie grant agreement No 101180586, the European Research Council (ERC) under the European Union’s Horizon 2020 research and innovation programme (grant agreement No 101001470) and the project National Institute of virology and bacteriology (Programme EXCELES, ID Project No. LX22NPO5103) - Funded by the European Union - Next Generation EU. Computational resources were provided by the Ministry of Education, Youth and Sports of the Czech Republic through the e-INFRA CZ (ID:90254).

## Author Contributions

Both W.M. and R.V. designed the research and wrote/edited the manuscript. W.M. performed the research and analysis.

## 5 Supplementary Information

### Forcefield

All simulations employed the CALVADOS2 coarse-grained force field[23], which represents each amino acid as a single bead centred at the C*α* position with implicit solvent. The total potential energy is

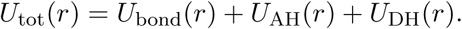

Adjacent beads are connected by a harmonic bond,

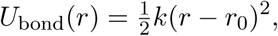

with *r*_0_ = 0.38 nm and *k* = 8368 kJ mol*^−^*^1^ nm*^−^*^2^ (2000 kcal mol*^−^*^1^ nm*^−^*^2^). Non-bonded interactions between residues separated by one bond are excluded.

Short-range interactions use a truncated and shifted Ashbaugh–Hatch potential,

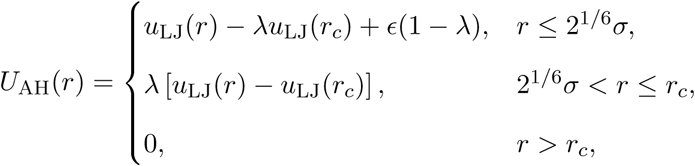

with

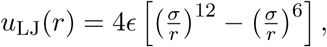

*ɛ* = 0.8368 kJ mol*^−^*^1^ and *r_c_* = 2.0 nm. Pair parameters are combined arithmetically, *σ_ij_* = (*σ_i_*+*σ_j_*)*/*2 and *λ_ij_* = (*λ_i_*+*λ_j_*)*/*2, using the CALVADOS2 residue-specific size (*σ*) and stickiness (*λ*) tabulations.

Salt-screened electrostatics are modelled with a Debye–Hückel potential truncated and shifted at 4.0 nm,

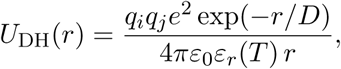

where *q_i_* are residue charges. Chain termini are treated as charged: +1 is added to the first residue and *−*1 to the last of every chain (with the corresponding terminal masses, +2 and +16 Da). The temperature-dependent dielectric constant is

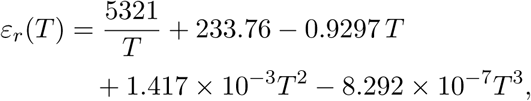

and the Debye length follows from the ionic strength c*_s_* = 0.15 M and the Bjerrum length B as 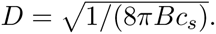

### 5.1 Demixing Normalization

Consider the limiting case in which both species condense independently to the same dense concentration *ρ*, so that every voxel is in one of three pure states: pure IDR-1 at (*ρ,* 0), pure IDR-2 at (0*, ρ*), or empty at (0, 0). If the dense phase occupies a volume fraction *a* of the box, the global composition constraint fixes the voxel fractions of the two dense states at *aϕ*_1_ and *aϕ*_2_. Projecting these three states onto the fixed basis and writing *n* = |(*ϕ*_1_*, ϕ*_2_)| gives

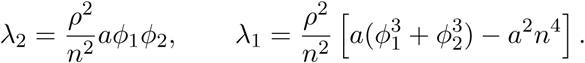

In a slab geometry the dense phase occupies a small fraction of the box, and in the limit *a →* 0 the quadratic term is negligible, so

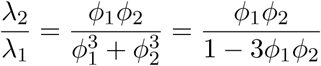

using 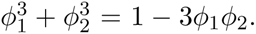 The perpendicular variance is therefore the smaller of the two under complete segregation, and substituting into *D* = 2 min(*λ*_1_*, λ*_2_)*/*(*λ*_1_ + *λ*_2_) gives

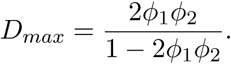

Because *D_max_*is derived in the *a →* 0 limit, a fully segregated system can marginally exceed it; the reported ratio (*D/D_max_*) is therefore capped at 1.

### 5.2 Comparing the Demixing Index

Here, we compare our demixing index against the mixing parameter recently introduced for polymer systems by Chen and Jacobs[27]. This method calculates the cosine similarity between the local composition (*ϕ⃗*) and the target composition (*ϕ⃗_a_*) over the one-dimensional density profile.

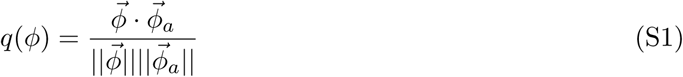

We set (*ϕ⃗_a_*) to the concentration ratio of each system. The *z* axis was discretised into 100 bins and time-averaged number densities *ρ*_1_(*z*) and *ρ*_2_(*z*) computed over the production run after 1 *µ*s equilibration. Only systems with a stable heterotypic condensate (Δ*G_ij_ < −*3 *k_B_T*) were included in the comparison. Agreement between *⟨q⟩_den_* and *D* was quantified by the square of the Pearson correlation coefficient, computed separately for three composition classes: equal (35–65% IDR-1), unequal (15–35% and 65–85%), and minority (below 15% and above 85%).

There is excellent agreement between the methods for equal residue composition (50:50 R^2^ = 0.97). However, the cosine-similarity approach requires an explicit dense-phase boundary over which to average. When the ratio deviates from (50:50) this dependency degrades the correlation with our metric (R^2^ < 0.5), where the dense phase of the minority component is often narrow, as exemplified in Supplementary Figure S6. In these regimes, our demixing index proves more robust, as it uses the variance across the entire concentration space without the need to define surface boundaries. While significantly larger simulations might mitigate the boundary artifacts affecting the cosine metric, our approach remains effective even for the system sizes typical of high-throughput studies.

**Table S1.** GIN clusters of intrinsically disordered regions (IDRs), recreated from Ruff et al.[37].

| Cluster | Description | General Category |
| --- | --- | --- |
| 0 | Blocks of G & polar residues | Polar |
| 1 | Blocks of P & polar residues | Polar |
| 2 | Blocks of negative P & G residues | Polar |
| 3 | Small negative blocks | Charged |
| 4 | Weak negative charge high N fraction | Polar |
| 5 | Blocks of positive negative & P residues | Polar |
| 6 | S patches | Polar |
| 7 | D/E-tracts | Charged |
| 8 | Blocks of negative & A residues | Charged |
| 9 | Blocks of positive & negative residues | Charged |
| 10 | Well-mixed hydrophobics enriched in M | Hydrophobic |
| 11 | Q-tracts | Polar |
| 12 | Blocks of positive negative & polar residues | Polar |
| 13 | Blocks of negative P & polar residues | Polar |
| 14 | Blocks of positive & P residues | Polar |
| 15 | T patches | Polar |
| 16 | Blocks of polar residues | Polar |
| 17 | Weak positive charge | Charged |
| 18 | Large negative blocks with positive blocks | Charged |
| 19 | High negative fraction specifically Es | Charged |
| 20 | A blocks | Hydrophobic |
| 21 | Blocks of P G A & polar residues | Polar |
| 22 | Well-mixed P and G | Polar |
| 23 | K blocks | Charged |
| 24 | Weak negative charge | Charged |
| 25 | Blocks of positive residues | Charged |
| 26 | R patches | Charged |
| 27 | P patches | Polar |
| 28 | High aromatic fraction specifically Ys | Hydrophobic |
| 29 | High R fraction | Charged |

**Figure S1.**
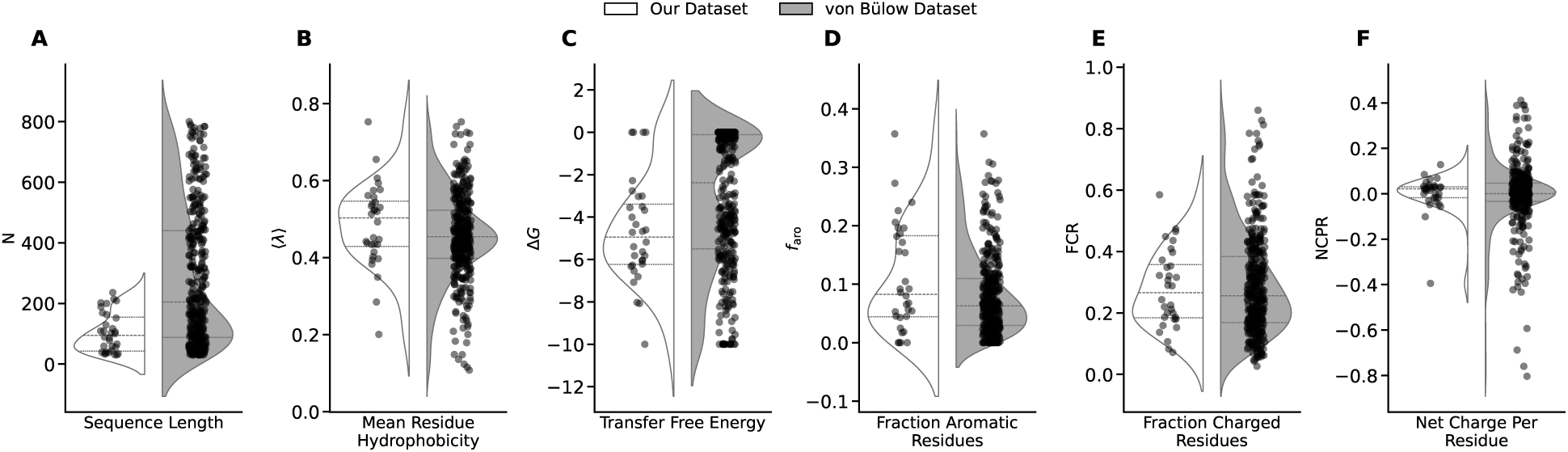
Distributions of key sequence features across selected IDRs (white/left) compared to the full von Bülow et al. dataset (grey/right)[33]. **(A)** Sequence length (N, amino acids). Our subset median (80 residues) is much lower than the dataset, but more closely matches the human IDR proteome[15]. **(B)** Mean residue hydrophobicity (⟨λ⟩), where values range from 0 (hydrophilic) to 1 (hydrophobic). **(C)** Transfer free energy (Δ*G*, *k_B_T*). More negative values indicate stronger homotypic phase separation propensity; Δ*G* ≈ 0 indicates no condensate formation. **(D)** Fraction of aromatic residues (*F*_aro_). **(E)** Fraction of charged residues (FCR). **(F)** Net charge per residue (NCPR).

**Figure S2.**
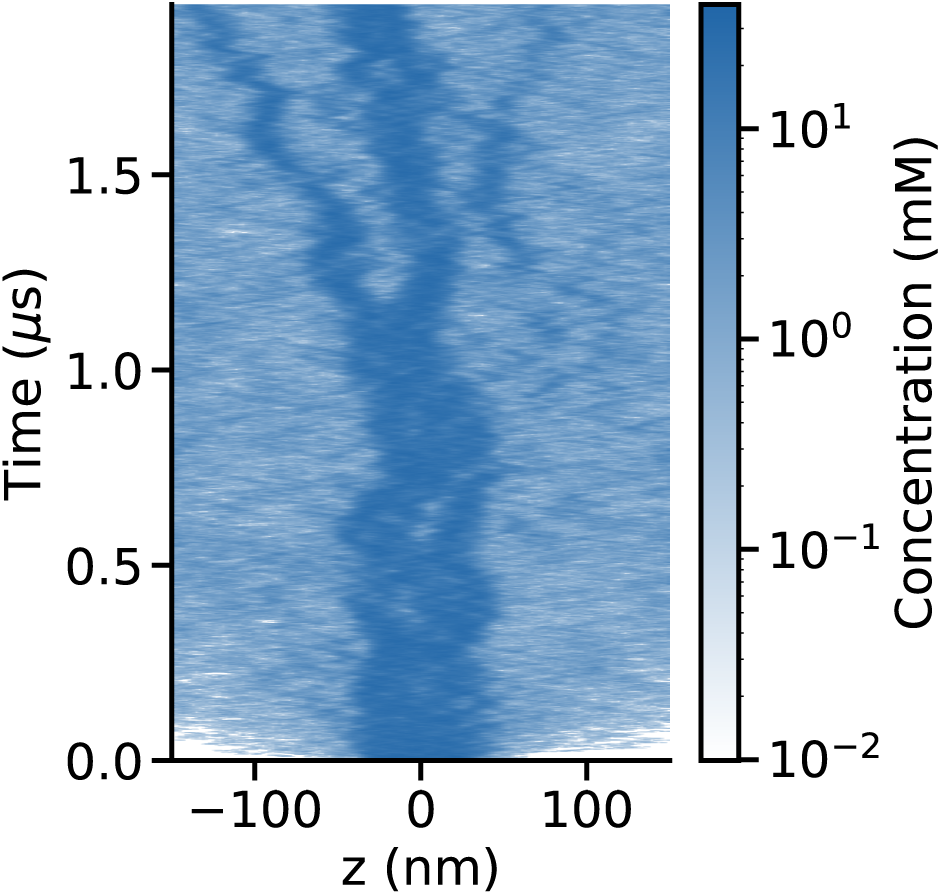
Concentration of the hnRNPA1 low-complexity domain along the elongated axis of the simulation cell as a function of time, at 311 K. Each row is the density profile of one frame, colored on a logarithmic scale, with z measured relative to the center of the cell. Rather than a single stable slab, two or more dense regions coexist and diffuse independently, occasionally coalescing and separating again over the course of the trajectory.

**Figure S3.**
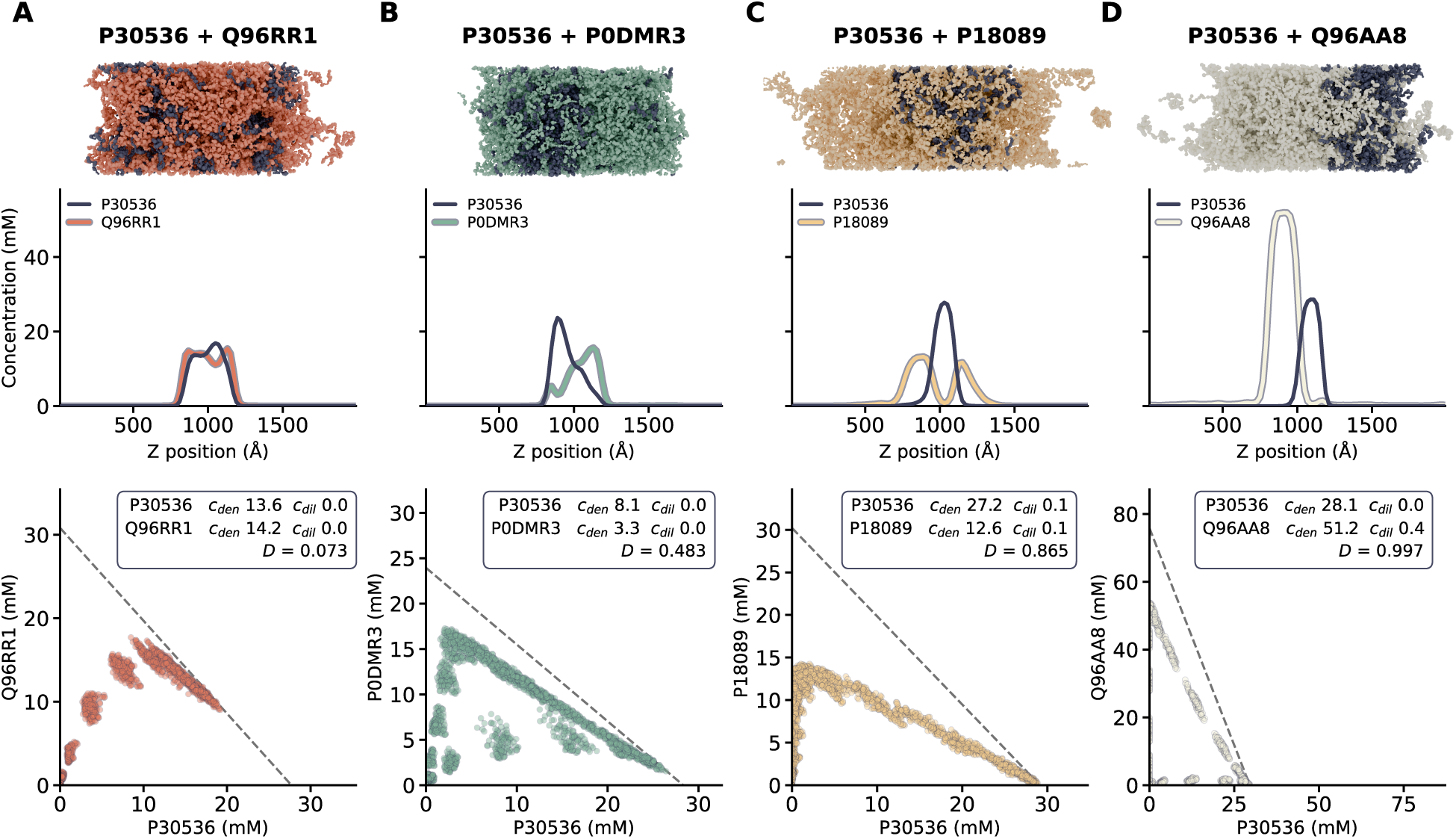
Condensate architectures of P30536 mixtures at (50:50) composition. Four mixtures of P30536 (dark blue) with partners of increasing demixing index. **Top:** final simulation frame. **Middle:** time-averaged concentration of each species along the long axis of the box. **Bottom:** concentration space, one point per voxel, with the dense- and dilute-phase concentrations of each component and the demixing index inset; the dashed line marks a constant total residue density, which is displaced from the diagonal because converting to mM divides each species by its own chain length. Concentrations are in mM throughout. **(A)** P30536 and Q96RR1 mix almost homogeneously (D = 0.073) and the two profiles overlap across the dense phase. However we can still view preference for P30536 self-affinity. This is captured in the concentration space, but the one dimensional profile is not uniform. **(B)** P0DMR3 (D = 0.483) partially mixes with P30536, but is highly enriched at the surface of the condensate **(C)** P18089 (D = 0.865) flanks P30536 on both sides, creating a core-shell structure. In the concentration space, only two interfaces are present (P18089:P30536 and P18089:Dilute). **(D)** Q96AA8 (D = 0.997) segregates into a separate condensate that shares a single interface with P30536. Three separate interfaces are shown in the concentration space, with two lying along the axes, and the third connecting the two dense phases. None of these architectures is radially symmetric about a condensate center, nor is there a homogeneous concentration of either protein throughout.

**Figure S4.**
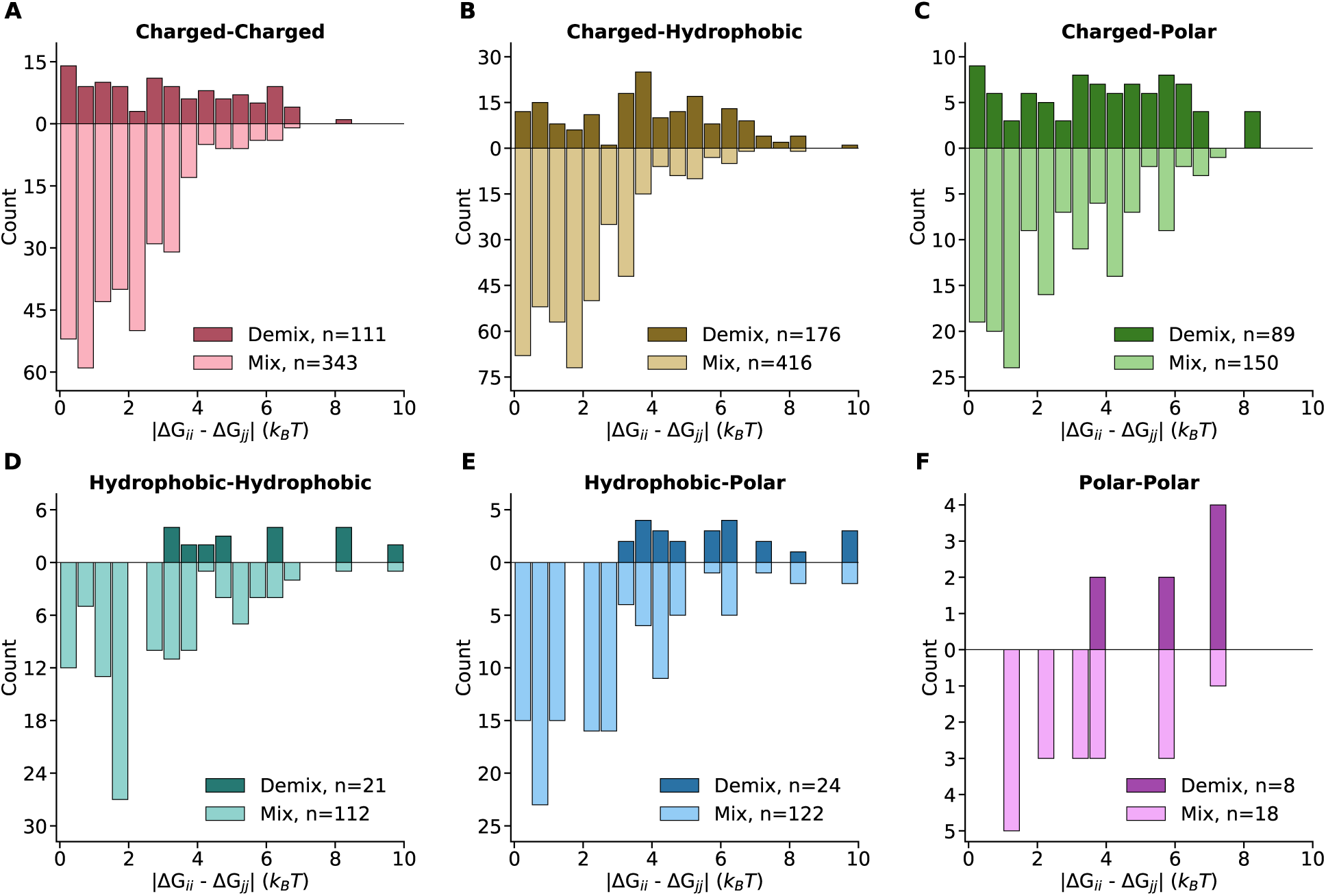
Demixing behavior by interaction type. Breakdown of Figure 3D by general interaction category: the distribution of |Δ*G_ii_ −* Δ*G_jj_|* for pairs classified as well-mixed (*D <* 0.2 and Δ*G_ij_ < −*3 *k_B_T*) or demixed (*D ≥* 0.2 and at least one homotypic Δ*G < −*3 *k_B_T*), using the same criteria as Figure 3C–D. **(A)** Charge-Charge, **(B)** Charge-Hydrophobic, **(C)** Charge-Polar, **(D)** Hydrophobic-Hydrophobic, **(E)** Hydrophobic-Polar, **(F)** Polar-Polar.

**Figure S5.**
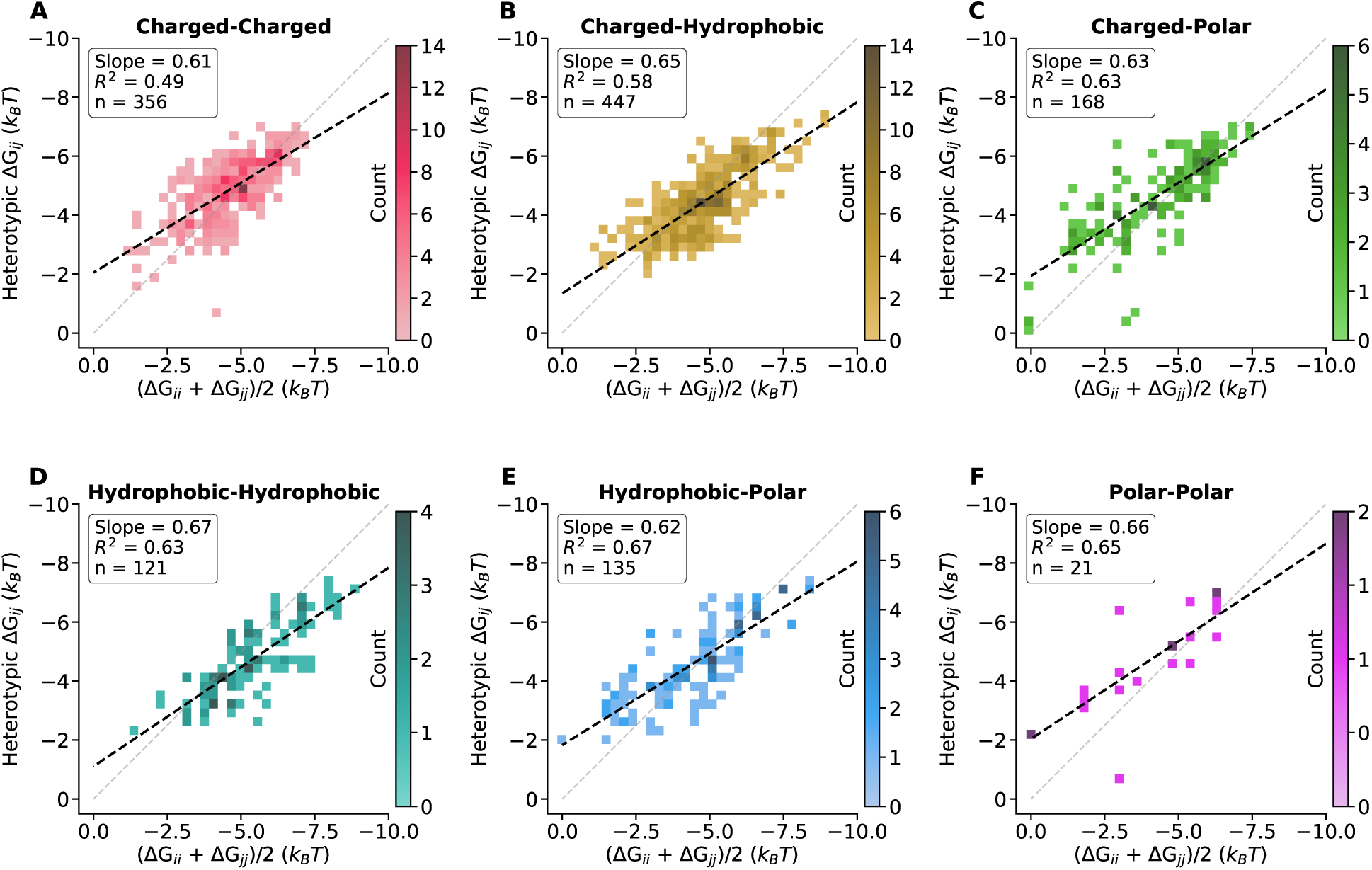
Heterotypic condensate stability stratified by interaction type. Analysis of heterotypic Δ*G_ij_* versus the average homotypic phase separation propensity ((Δ*G_ii_* +Δ*G_jj_*)/2) for each general interaction category. **(A)** Charge-Charge, **(B)** Charge-Hydrophobic, **(C)** Charge-Polar, **(D)** Hydrophobic-Hydrophobic, **(E)** Hydrophobic-Polar, **(F)** Polar-Polar. Each panel shows a 2D histogram with a linear regression fit (black dashed line) and the identity line (gray dashed line) for reference. Inset statistics report the slope, *R*^2^, and sample size for each interaction type. Slopes near 1.0 indicate that heterotypic stability closely tracks the average of homotypic values, while deviations suggest interaction-specific effects.

**Figure S6.**
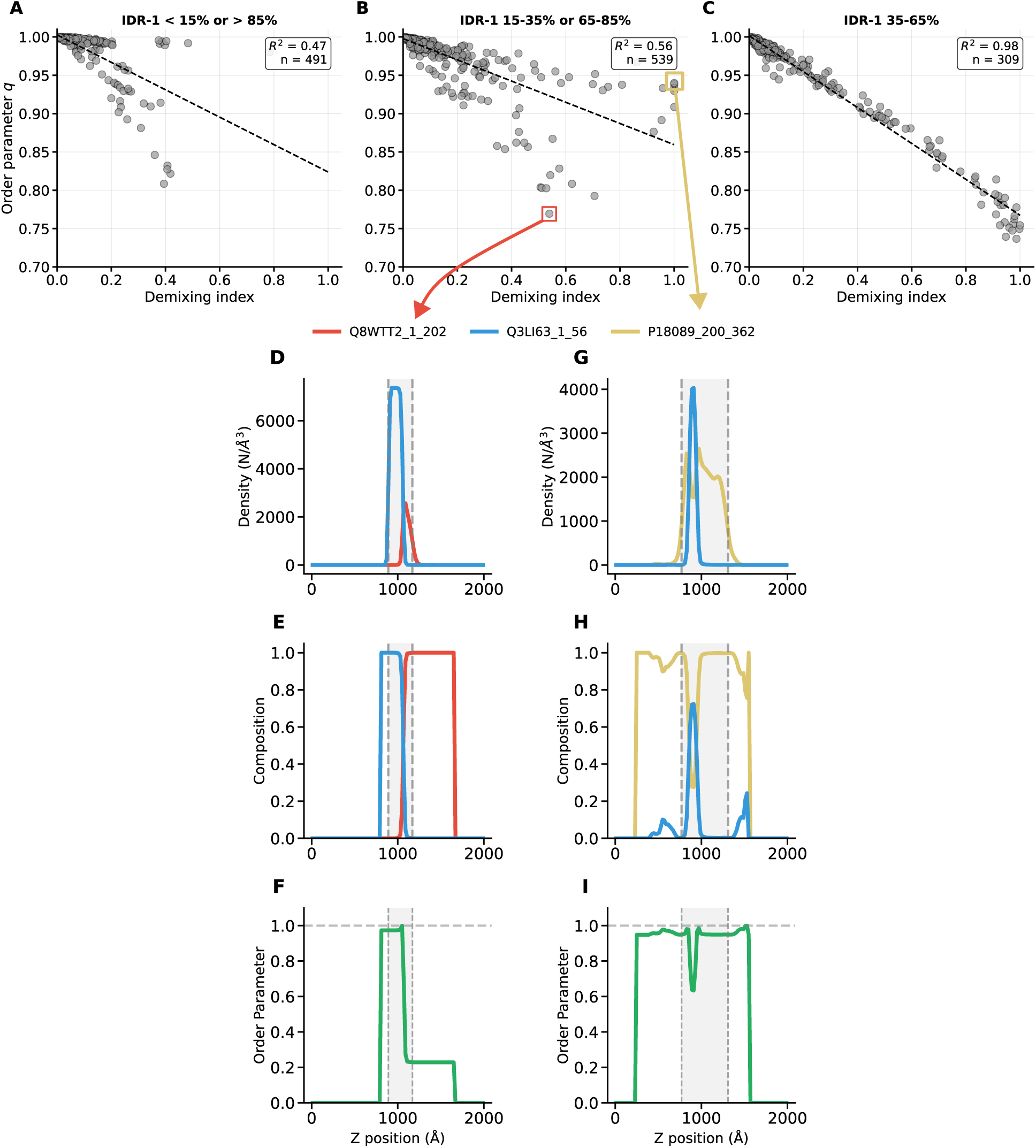
Comparison of Demixing Index with Local Order Parameter(A-C) Correlation between the Demixing Index (D) and the mixing order parameter (q, cosine similarity) across three compositional regimes with respect to IDR-1: **(A)** *<* 15% or *>* 85% **(B)** 15–35% or 65–85%, and **(C)** Equal (50%). The metrics show strong agreement at (50:50) compositions, but diverge significantly at asymmetric compositions. **(D-I)** Detailed 1-dimensional density profiles of two systems highlighting the sensitivity of the order parameter to phase boundary definitions. The left column (D-F) shows a mixture of Q8WTT2 (red - 19%) and Q3LI63 (blue - 81%), while the right column (G-I) shows a mixture of Q3LI63 (25%) and P18089 (yellow - 75%). **(D, G)** Density profiles along the Z-axis for the two IDRs. The gray-shaded region indicates the dense phase as identified by a density cutoff. **(E, H)** Local composition fractions along the Z-axis. **(F, I)** The local order parameter is calculated along the Z-axis. The dashed line represents perfect mixing (q=1). In asymmetric mixtures (Left), the minority component often localizes at the interface or forms a gradient, making the calculation of an average order parameter highly sensitive to the precise definition of the dense-phase boundaries (gray region).36Our Demixing Index circumvents this by calculating variance across the entire concentration space, independent of explicit spatial boundaries.

## References

[1] Daniel S. W. Lee, Amy R. Strom, and Clifford P. Brangwynne. The mechanobiology of nuclear phase separation. APL Bioengineering, 6(2), 2022. doi: 10.1063/5.0083286.

[2] Alex S. Holehouse and Birthe B. Kragelund. The molecular basis for cellular function of intrinsically disordered protein regions. Nature Reviews Molecular Cell Biology, 25(3):187– 211, 2024. doi: 10.1038/s41580-023-00673-0.

[3] Andrew G. T. Pyo, Yaojun Zhang, and Ned S. Wingreen. Proximity to criticality predicts surface properties of biomolecular condensates. Proceedings of the National Academy of Sciences, 120(23):e2220014120, 2023. doi: 10.1073/pnas.2220014120.

[4] Kathleen A. Burke, Abigail M. Janke, Christy L. Rhine, and Nicolas L. Fawzi. Residue-by-residue view of in vitro FUS granules that bind the C-terminal domain of RNA polymerase II. Molecular Cell, 60(2):231–241, 2015. doi: 10.1016/j.molcel.2015.09.006. URL https://doi.org/10.1016/j.molcel.2015.09.006.

[5] Nathaniel Hess and Jerelle A. Joseph. Structured protein domains enter the spotlight: modulators of biomolecular condensate form and function. Trends in Biochemical Sciences, 50(3): 206–223, 2025. doi: 10.1016/j.tibs.2024.12.008. URL https://doi.org/10.1016/j.tibs.2024.12.008.

[6] V. Pandey, T. Hosokawa, Y. Hayashi, and H. Urakubo. Multiphasic protein condensation governed by shape and valency. Cell Reports, 44(4):115504, 2025. doi: 10.1016/j.celrep.2025.115504. URL https://doi.org/10.1016/j.celrep.2025.115504.

[7] Rachel M. Welles, Kandarp A. Sojitra, Mikael V. Garabedian, Boao Xia, Wentao Wang, Muyang Guan, Roshan M. Regy, Elizabeth R. Gallagher, Daniel A. Hammer, Jeetain Mittal, and Matthew C. Good. Determinants that enable disordered protein assembly into discrete condensed phases. Nature Chemistry, 16(7):1062–1072, 2024. doi: 10.1038/s41557-023-01423-7.

[8] William M. Jacobs. Self-Assembly of Biomolecular Condensates with Shared Components. Physical Review Letters, 126(25):258101, 2021. doi: 10.1103/PhysRevLett.126.258101.

[9] M. Biesaga, M. Frigolé-Vivas, and X. Salvatella. Intrinsically disordered proteins and biomolecular condensates as drug targets. Current Opinion in Chemical Biology, 62:90–100, 2021. doi: 10.1016/j.cbpa.2021.02.009.

[10] J. E. Bradner, D. Hnisz, and R. A. Young. Transcriptional addiction in cancer. Cell, 168(4): 629–643, 2017. doi: 10.1016/j.cell.2016.12.013.

[11] L. Jiang and Y. Kang. Biomolecular condensates: A new lens on cancer biology. Biochimica et Biophysica Acta (BBA) - Reviews on Cancer, 1880(1):189245, 2025. doi: 10.1016/j.bbcan.2024.189245.

[12] A. Patel, H. O. Lee, L. Jawerth, S. Maharana, M. Jahnel, M. Y. Hein, S. Stoynov, J. Mahamid, S. Saha, T. M. Franzmann, A. Pozniakovski, I. Poser, N. Maghelli, L. A. Royer, M. Weigert, E. W. Myers, S. Grill, D. Drechsel, A. A. Hyman, and S. Alberti. A liquid-to-solid phase transition of the ALS protein FUS accelerated by disease mutation. Cell, 162(5):1066–1077, 2015. doi: 10.1016/j.cell.2015.07.047.

[13] K. Tsuji, H. Kawata, T. Kamiakito, T. Nakaya, and A. Tanaka. RNA-binding protein 14 promotes phase separation to sustain prostate specific antigen expression under androgen deprivation in human prostate cancer. The Journal of Steroid Biochemistry and Molecular Biology, 235:106407, 2023. doi: 10.1016/j.jsbmb.2023.106407.

[14] Savannah E. LaBuda, Russell R. Broaddus, and Andrew B. Gladden. ARID1A: Gene, protein, and function in endometrial cancer. Oncogene, 44(36):3273–3283, 2025. doi: 10.1038/s41388-025-03539-1. URL https://www.nature.com/articles/s41388-025-03539-1.

[15] Giulio Tesei, Anna Ida Trolle, Nicolas Jonsson, Johannes Betz, Frederik E. Knudsen, Francesco Pesce, Kristoffer E. Johansson, and Kresten Lindorff-Larsen. Conformational ensembles of the human intrinsically disordered proteome. Nature, 626(8000):897–904, 2024. doi: 10.1038/s41586-023-07004-5.

[16] Kathryn Tunyasuvunakool, Jonas Adler, Zachary Wu, Tim Green, Michal Zielinski, Augustin Žídek, Alex Bridgland, Andrew Cowie, Clemens Meyer, Agata Laydon, Sameer Velankar, Gerard J. Kleywegt, Alex Bateman, Richard Evans, Alexander Pritzel, Michael Figurnov, Olaf Ronneberger, Russ Bates, Simon A. A. Kohl, Anna Potapenko, Andrew J. Ballard, Bernardino Romera-Paredes, Stanislav Nikolov, Rishub Jain, Ellen Clancy, David Reiman, Stig Petersen, Andrew W. Senior, Koray Kavukcuoglu, Ewan Birney, Pushmeet Kohli, John Jumper, and Demis Hassabis. Highly accurate protein structure prediction for the human proteome. Nature, 596(7873):590–596, 2021. doi: 10.1038/s41586-021-03828-1. URL https://doi.org/10.1038/s41586-021-03828-1.

[17] Daoyuan Qian, Timothy J. Welsh, Nadia A. Erkamp, Seema Qamar, Jonathon Nixon-Abell, Georg Krainer, Peter St. George-Hyslop, Thomas C. T. Michaels, and Tuomas P. J. Knowles. Tie-line analysis reveals interactions driving heteromolecular condensate formation. Physical Review X, 12(4):041038, 2022. doi: 10.1103/PhysRevX.12.041038. URL https://link.aps.org/doi/10.1103/PhysRevX.12.041038.

[18] Kerstin Dörner, Michelle Jennifer Gut, Daan Overwijn, Fan Cao, Matej Siketanc, Stephanie Heinrich, Nicole Beuret, Justin Meyer, Timothy Sharpe, Kresten Lindorff-Larsen, and Maria Hondele. Fluorescent protein and peptide tags alter condensate formation and dynamics in vivo and in vitro. EMBO Reports, 27(1):89–121, 2026. doi: 10.1038/s44319-025-00626-y. URL https://doi.org/10.1038/s44319-025-00626-y.

[19] Gregory L. Dignon, Wenwei Zheng, Young C. Kim, and Jeetain Mittal. Temperature-controlled liquid–liquid phase separation of disordered proteins. ACS Central Science, 5(5):821–830, 2019. doi: 10.1021/acscentsci.9b00102. URL https://doi.org/10.1021/acscentsci.9b00102.

[20] Gregory L. Dignon, Wenwei Zheng, Robert B. Best, Young C. Kim, and Jeetain Mittal. Relation between single-molecule properties and phase behavior of intrinsically disordered proteins. Proceedings of the National Academy of Sciences, 115(40):9929–9934, 2018. doi: 10.1073/pnas.1804177115. URL https://www.pnas.org/doi/abs/10.1073/pnas.1804177115.

[21] Alejandro Feito, Ignacio Sanchez-Burgos, Ignacio Tejero, Eduardo Sanz, Antonio Rey, Rosana Collepardo-Guevara, Andrés R. Tejedor, and Jorge R. Espinosa. Benchmarking residue-resolution protein coarse-grained models for simulations of biomolecular condensates. PLOS Computational Biology, 21(1):1–31, 2025. doi: 10.1371/journal.pcbi.1012737. URL https://doi.org/10.1371/journal.pcbi.1012737.

[22] Andrés R. Tejedor, Anne Aguirre Gonzalez, M. Julia Maristany, Pin Yu Chew, Kieran Russell, Jorge Ramirez, Jorge R. Espinosa, and Rosana Collepardo-Guevara. Chemically Informed Coarse-Graining of Electrostatic Forces in Charge-Rich Biomolecular Condensates. ACS Central Science, 11(2):302–321, 2025. doi: 10.1021/acscentsci.4c01617. URL https://doi.org/10.1021/acscentsci.4c01617.

[23] Giulio Tesei and Kresten Lindorff-Larsen. Improved predictions of phase behaviour of intrinsically disordered proteins by tuning the interaction range. Open Research Europe, 2:94, 2023. doi: 10.12688/openreseurope.14967.2.

[24] Mina Farag, Wade M. Borcherds, Anne Bremer, Tanja Mittag, and Rohit V. Pappu. Phase separation of protein mixtures is driven by the interplay of homotypic and heterotypic interactions. Nature Communications, 14(1):5527, 2023. doi: 10.1038/s41467-023-41274-x. URL https://doi.org/10.1038/s41467-023-41274-x.

[25] Kyosuke Adachi and Kyogo Kawaguchi. Predicting Heteropolymer Interactions: Demixing and Hypermixing of Disordered Protein Sequences. Physical Review X, 14(3):031011, 2024. doi: 10.1103/physrevx.14.031011.

[26] Pin Yu Chew, Jerelle A. Joseph, Rosana Collepardo-Guevara, and Aleks Reinhardt. Thermodynamic origins of two-component multiphase condensates of proteins. Chemical Science, 14(7):1820–1836, 2023. doi: 10.1039/D2SC05873A. URL http://dx.doi.org/10.1039/D2SC05873A.

[27] Fan Chen and William M. Jacobs. Emergence of multiphase condensates from a limited set of chemical building blocks. Journal of Chemical Theory and Computation, 20(15):6881–6889, 2024. doi: 10.1021/acs.jctc.4c00323. URL https://doi.org/10.1021/acs.jctc.4c00323.

[28] Tamas Lazar, Acadia Connor, Charles F. DeLisle, Virginia Burger, and Peter Tompa. Targeting protein disorder: The next hurdle in drug discovery. Nature Reviews Drug Discovery, 24: 743–763, 2025. doi: 10.1038/s41573-025-01220-6.

[29] Caixuan Liu, Kejia Wu, Hojun Choi, Hannah L. Han, Xueli Zhang, Joseph L. Watson, Green Ahn, Jason Z. Zhang, Sara Shijo, Lydia L. Good, Charlotte M. Fischer, Asim K. Bera, Alex Kang, Evans Brackenbrough, Brian Coventry, Derrick R. Hick, Seema Qamar, Xinting Li, Justin Decarreau, Stacey R. Gerben, Wei Yang, Inna Goreshnik, Dionne Vafeados, Xinru Wang, Mila Lamb, Analisa Murray, Sebastian Kenny, Magnus S. Bauer, Andrew N. Hoofnagle, Ping Zhu, Tuomas P. J. Knowles, and David Baker. Diffusing protein binders to intrinsically disordered proteins. Nature, 644(8077):809–817, 2025. doi: 10.1038/s41586-025-09248-9. URL https://doi.org/10.1038/s41586-025-09248-9.

[30] Lev D. Gelb and Erich A. Müller. Location of phase equilibria by temperature-quench molecular dynamics simulations. Fluid Phase Equilibria, 203(1-2):1–14, 2002. doi: 10.1016/s0378-3812(02)00174-7.

[31] M Rovere, D W Heermann, and K Binder. The gas-liquid transition of the two-dimensional Lennard-Jones fluid. Journal of Physics: Condensed Matter, 2(33):7009–7032, 1990. doi: 10.1088/0953-8984/2/33/013.

[32] H. Flyvbjerg and H. G. Petersen. Error estimates on averages of correlated data. The Journal of Chemical Physics, 91(1):461–466, 1989. doi: 10.1063/1.457480. URL https://doi.org/10.1063/1.457480.

[33] S. von Bülow, G. Tesei, F.K. Zaidi, T. Mittag, and K. Lindorff-Larsen. Prediction of phase-separation propensities of disordered proteins from sequence. Proceedings of the National Academy of Sciences, 122(13):e2417920122, 2025. doi: 10.1073/pnas.2417920122. URL https://doi.org/10.1073/pnas.2417920122.

[34] Zakarya Benayad, Sören von Bülow, Lukas S Stelzl, and Gerhard Hummer. Simulation of FUS protein condensates with an adapted Coarse-Grained model. Journal of Chemical Theory and Computation, 17(1):525–537, 2021. doi: 10.1021/acs.jctc.0c01064.

[35] Gaurav Mitra, Souradeep Ghosh, Kiersten M. Ruff, Ruoyao Zhang, Gaurav Chauhan, and Rohit V. Pappu. Distinguishing near- versus off-critical phase behaviors of intrinsically disordered proteins. Reports on Progress in Physics, 89(6):068101, 2026. doi: 10.1088/1361-6633/ae70d6.

[36] Andrew Z. Lin, Kiersten M. Ruff, Furqan Dar, Ameya Jalihal, Matthew R. King, Jared M. Lalmansingh, Ammon E. Posey, Nadia A. Erkamp, Ian Seim, Amy S. Gladfelter, and Rohit V. Pappu. Dynamical control enables the formation of demixed biomolecular condensates. Nature Communications, 14(1):7678, 2023. doi: 10.1038/s41467-023-43489-4. URL https://doi.org/10.1038/s41467-023-43489-4.

[37] Kiersten M. Ruff, Matthew R. King, Alexander W. Ying, Vicky Liu, Avnika Pant, Whitney E. Lieberman, Min Kyung Shinn, Xiaolei Su, Cigall Kadoch, and Rohit V. Pappu. Molecular grammars of predicted intrinsically disordered regions that span the human proteome. Cell, 189(1):323–342.e17, 2026. doi: 10.1016/j.cell.2025.10.019. URL https://doi.org/10.1016/j.cell.2025.10.019.

[38] Simon M. Lichtinger, Adiran Garaizar, Rosana Collepardo-Guevara, and Aleks Reinhardt. Targeted modulation of protein liquid–liquid phase separation by evolution of amino-acid sequence. PLOS Computational Biology, 17(8):1–28, 2021. doi: 10.1371/journal.pcbi.1009328. URL https://doi.org/10.1371/journal.pcbi.1009328.

[39] Jie Wang, Jeong-Mo Choi, Alex S. Holehouse, Hyun O. Lee, Xiaojie Zhang, Marcus Jahnel, Shovamayee Maharana, Régis Lemaitre, Andrei Pozniakovsky, David Drechsel, Ina Poser, Rohit V. Pappu, Simon Alberti, and Anthony A. Hyman. A molecular grammar governing the driving forces for phase separation of prion-like RNA binding proteins. Cell, 174(3): 688–699.e16, 2018. doi: 10.1016/j.cell.2018.06.006. URL https://www.sciencedirect.com/science/article/pii/S0092867418307311.

[40] Gaofeng Pei, Xinxin Wang, Xuebo Quan, Danqian Geng, Zhuo Chen, Weifan Xu, Kai Huang, Tingting Li, and Pilong Li. Opposing roles of serine and charge in IDR condensate miscibility. Nature Chemical Biology, 22(8):1274–1285, 2026. doi: 10.1038/s41589-026-02251-9. URL https://doi.org/10.1038/s41589-026-02251-9.

[41] Yong Ryoul Kim, Jaegeon Joo, Hee Jung Lee, Chaelim Kim, Ju-Chan Park, Young Suk Yu, Chang Rok Kim, Do Hui Lee, Joowon Cha, Hyemin Kwon, Kimberley M. Hanssen, Thomas G. P. Grünewald, Murim Choi, Ilkyu Han, Sangsu Bae, Inkyung Jung, Yongdae Shin, and Sung Hee Baek. Prion-like domain mediated phase separation of ARID1A promotes oncogenic potential of Ewing’s sarcoma. Nature Communications, 15(1):6569, 2024. doi: 10.1038/s41467-024-51050-0. URL https://doi.org/10.1038/s41467-024-51050-0.

[42] Liling Wan, Shasha Chong, Fan Xuan, Angela Liang, Xiaodong Cui, Leah Gates, Thomas S. Carroll, Yuanyuan Li, Lijuan Feng, Guochao Chen, Shu-Ping Wang, Michael V. Ortiz, Sara K. Daley, Xiaolu Wang, Hongwen Xuan, Alex Kentsis, Tom W. Muir, Robert G. Roeder, Haitao Li, Wei Li, Robert Tjian, Hong Wen, and C. David Allis. Impaired cell fate through gain-of-function mutations in a chromatin reader. Nature, 577(7788):121–126, 2020. doi: 10.1038/s41586-019-1842-7. URL https://www.nature.com/articles/s41586-019-1842-7.

[43] Y. Dai, Z. Zhou, W. Yu, Y. Ma, K. Kim, N. Rivera, J. Mohammed, E. Lantelme, H. Hsu-Kim, A. Chilkoti, and L. You. Biomolecular condensates regulate cellular electrochemical equilibria. Cell, 187(21):5951–5966.e18, 2024. doi: 10.1016/j.cell.2024.08.018.

[44] A. S. Holehouse, R. K. Das, J. N. Ahad, M. O. G. Richardson, and R. V. Pappu. CIDER: Resources to analyze sequence-ensemble relationships of intrinsically disordered proteins. Biophysical Journal, 112(1):16–21, 2017. doi: 10.1016/j.bpj.2016.11.3200. URL https://doi.org/10.1016/j.bpj.2016.11.3200.

[45] Heankel Lyons, Reshma T. Veettil, Prashant Pradhan, Christy Fornero, Nancy De La Cruz, Keiichi Ito, Mikayla Eppert, Robert G. Roeder, and Benjamin R. Sabari. Functional partitioning of transcriptional regulators by patterned charge blocks. Cell, 186(2):327–345.e28, 2023. doi: 10.1016/j.cell.2022.12.013. URL https://www.sciencedirect.com/science/article/pii/S0092867422015264.

[46] Atar Gilat, Benjamin Dubrueil, and Emmanuel D. Levy. Mapping interactions between disordered regions reveals promiscuity in biomolecular condensate formation. bioRxiv, 2023. doi: 10.1101/2023.07.04.547715. URL https://www.biorxiv.org/content/early/2023/07/04/2023.07.04.547715.

[47] Daoyuan Qian, Hannes Ausserwoger, Tomas Sneideris, Mina Farag, Rohit V. Pappu, and Tuomas P. J. Knowles. Dominance analysis to assess solute contributions to multicomponent phase equilibria. Proceedings of the National Academy of Sciences, 121(33):e2407453121, 2024. doi: 10.1073/pnas.2407453121. URL https://www.pnas.org/doi/abs/10.1073/pnas.2407453121.

[48] Brian D. Freibaum, James Messing, Peiguo Yang, Hong Joo Kim, and J. Paul Taylor. High-fidelity reconstitution of stress granules and nucleoli in mammalian cellular lysate. Journal of Cell Biology, 220(3):e202009079, 2021. doi: 10.1083/jcb.202009079. URL https://doi.org/10.1083/jcb.202009079.

[49] Ikki Yasuda, Sören von Bülow, Giulio Tesei, Eiji Yamamoto, Kenji Yasuoka, and Kresten Lindorff-Larsen. Coarse-grained model of disordered RNA for simulations of biomolecular condensates. Journal of Chemical Theory and Computation, 21(5):2766–2779, 2025. doi: 10.1021/acs.jctc.4c01646. URL https://doi.org/10.1021/acs.jctc.4c01646.

[50] Jonathan E. Henninger, Ozgur Oksuz, Krishna Shrinivas, Ido Sagi, Gary LeRoy, Ming M. Zheng, J. Owen Andrews, Alicia V. Zamudio, Charalampos Lazaris, Nancy M. Hannett, Tong Ihn Lee, Phillip A. Sharp, Ibrahim I. Cissé, Arup K. Chakraborty, and Richard A. Young. RNA-mediated feedback control of transcriptional condensates. Cell, 184(1):207–225.e24, 2021. doi: 10.1016/j.cell.2020.11.030. URL https://www.sciencedirect.com/science/article/pii/S0092867420315671.

[51] Ikki Yasuda, Eiji Yamamoto, Kenji Yasuoka, and Kresten Lindorff-Larsen. Molecular interactions underlying selective partitioning of clients into MED1 and FUS condensates. bioRxiv, 2025. doi: 10.64898/2025.12.23.696191. URL https://www.biorxiv.org/content/early/2025/12/23/2025.12.23.696191.

[52] Alexander E. Conicella, Gregory L. Dignon, Gül H. Zerze, Hermann Broder Schmidt, Alexandra M. D’Ordine, Young C. Kim, Rajat Rohatgi, Yuna M. Ayala, Jeetain Mittal, and Nicolas L. Fawzi. TDP-43 α-helical structure tunes liquid–liquid phase separation and function. Proceedings of the National Academy of Sciences, 117(11):5883–5894, 2020. doi: 10.1073/pnas.1912055117. URL https://www.pnas.org/doi/abs/10.1073/pnas.1912055117.

[53] Peter Eastman, Jason Swails, John Chodera, Robert T. McGibbon, Andy Simmonett, Michael Sherman, Lee-Ping Wang, Rafal Wiewiora, Charlles Abreu, Saurabh Belsare, Yutong Zhao, Kyle Beauchamp, Chaya Stern, João Rodrigues, Anton Gorenko, Joy Ku, Sam Mikes, Marc Marí, Jing Huang, and Matthew Harrigan. openmm/openmm: OpenMM 8.2.0 (8.2.0), 2024. URL 10.5281/zenodo.14058389. Accessed: 2024-07-24.

